# A model of functionally buffered deleterious mutations can lead to signatures of positive selection

**DOI:** 10.1101/2022.02.14.480440

**Authors:** Runxi Shen, Miwa Wenzel, Philipp W Messer, Charles F. Aquadro

**Affiliations:** Cornell University

## Abstract

Selective pressures on DNA sequences often result in departures from neutral evolution that can be captured by the McDonald-Kreitman (MK) test. However, the nature of such selective forces often remains unknown to experimentalists. Amino acid fixations driven by natural selection in protein coding genes are commonly associated with a genetic arms race or changing biological purposes, leading to proteins with new functionality. Here, we evaluate the expectations of a buffering mechanism driving selective amino acids to fixation, which is motivated by an observed phenotypic rescue of otherwise deleterious nonsynonymous substitutions at *bag of marbles* (*bam*) and *Sex lethal* (*Sxl*). These two important genes in *Drosophila melanogaster* were shown to experience strong episodic bursts of natural selection potentially due to infections of the endosymbiotic bacteria *Wolbachia* observed among multiple *Drosophila* species. Using simulations to implement and evaluate the evolutionary dynamics of a *Wolbachia* buffering model, we demonstrate that selectively fixed amino acid replacements will occur, but that proportion of adaptive amino acid fixations and the statistical power of the MK test to detect the departure from an equilibrium neutral model are both significantly lower than seen for an arms race/change-in-function model that favors proteins with diversified amino acids.

## Introduction

Patterns of DNA sequence variation within and between species have been widely used to infer the evolutionary forces that have acted on genes and genomes. Over the past three decades, many statistical tests of a model of neutral evolution have been developed, with one of the most widely applied being the McDonald-Kreitman (MK) test (McDonald and Kreitman 1991). The basis of this test is a comparison of the ratios of nonsynonymous and synonymous fixed differences between species to those segregating as polymorphisms within species using a 2×2 contingency test (e.g., Fisher’s Exact Test). Synonymous variation is a proxy for neutral variation, and an excess of nonsynonymous fixed differences between species is typically interpreted as evidence that natural selection has accelerated the fixation of advantageous amino acid replacements. This pattern is often associated with natural selection fine tuning protein function, in response to a changing function and/or an intra- or inter-genomic arms race (e.g., adaptations by a genome to silence transposable elements and reciprocal adaptations between predator and prey, respectively) (McLaughlin and Malik 2017). Although the MK test has been found to have low power in detecting positive selection, particularly when applied to single genes (Akashi 1999; Zhai, et al. 2009), there are many empirical reports in the literature of significant departures in the direction of positive selection (e.g., Eyre-Walker 2006). The obvious question from the experimentalist’s perspective is what evolutionary mechanisms are driving such signatures of positive selection.

We have studied the population genetics of two *Drosophila* germline stem cell genes, *bag of marbles (bam)* and *Sex lethal (Sxl)*, that show strong evidence of episodic positive selection in several species (Bauer DuMont, et al. 2007; Flores, DuMont, et al. 2015; Bauer DuMont, et al. 2021). This positive selection has been proposed to be due to a change in gene function and/or an evolutionary arms race with the endosymbiont bacteria *Wolbachia* that genetically interacts with both genes and resides in the germarium where they function (Bauer DuMont, et al. 2007; Flores, Bubnell, et al. 2015; Flores, DuMont, et al. 2015; Bubnell, et al. 2021, Bauer DuMont, et al. 2021).

*Wolbachia* infections have been observed to be temporally dynamic in host populations, being lost at times and then regained (Richardson, et al. 2012; Turelli, et al. 2018; Meany, et al. 2019). We thus evaluate an alternative model based on standard population genetic theory and the observed rescue of fertility defects of four distinct amino acid replacements mutations of *bam* and *Sxl* in *D. melanogaster* (Flores, Bubnell, et al. 2015; Star and Cline, 2002). The *Wolbachia* rescue results suggest that during *Wolbachia* infection, slightly deleterious amino acid replacements might accumulate by drift in these (and potentially other) genes without significantly reducing *D. melanogaster* fitness. When *Wolbachia* is lost from the population, there could be positive selection for new nonsynonymous mutations that return the *bam* and *Sxl* protein sequences to their initial, and assumed optimal, functional state. We term this dynamic the Buffering model, as the effects of deleterious mutations are “buffered” during periods of infection by *Wolbachia*.

We implemented simulations to compare the evolutionary processes of our Buffering model with those from a classical arms race between the host germline gene and *Wolbachia.* For example, *Wolbachia* may manipulate *bam* and *Sxl* in a way counterproductive to the fitness of the fly. Arms race dynamics are expected to lead to positive selection favoring diversifying amino acids in these genes that result in *Drosophila’s* escape from the deleterious impact of *Wolbachia* on their fitness. Note that while we model this as an evolutionary arms race, the results of selection associated with a strong directional shift in function and thus amino acid sequence would be similar many ways.

We base our modeling on the *bam* gene alone because *Sxl* has a large and highly conserved RNA binding domain that limits the sites available for adaptive evolution in our simulations (Bauer DuMont et al. 2021). We demonstrate that buffering type of interaction does result in positive selection when we track the number of adaptive mutations that fix in the population during the simulations. However, application of the MK test to the resultant numbers of synonymous and nonsynonymous fixations and polymorphisms among sequences sampled at the end of the simulations detects only weak departures from neutrality consistent with positive selection. We also find that it is difficult to distinguish between the buffering or an arms race models based on properties of the amino acid substitutions observed, though an arms race model does lead to more adaptive fixations overtime.

## Materials and Methods

### Simulation model

The evolution of *bam* was simulated under a Wright-Fisher model using nucleotide-based simulation in SLiM 3.5 (Haller et al. 2019). We first inferred the ancestral DNA sequence of the exons in *bam* (1338 nucleotides) for *Drosophila melanogaster* and *D. simulans* using maximum likelihood with codeML v4.8 (Yang 2007). Briefly, alignments of seven *Drosophila* coding sequences were made using PRANK v.170427 (Loytynoja 2014), including sequences of *D. melanogaster, D. simulans, D. sechellia, D. yakuba, D. erecta, D. eugracilis, and D. pseudoobscura.* The alignment was input in codeML and an ancestral sequence was estimated by maximum likelihood for the common ancestor of *D. melanogaster* and *D. simulans*, using the other species’ sequences as outgroup references. The estimated common ancestral nucleotide sequence was then used in SLiM as the starting sequence for the entire population in each simulation. We simulated *bam* as a single contiguous exon, though in reality there are two short introns (61 and 64 bp). The effect of excluding these introns has a negligible impact on the rate of recombination.

The evolutionary parameters in the simulations were based on empirical estimates. In particular, we used an effective population size *(N_e_)* of 1e6 (Campos et al. 2017), an overall mutation probability Qz) of 2.8e-9 per nucleotide per generation, with a 2:1 transition:transversion rate (Keightley et al. 2009, Keightley et al. 2014), an average recombination rate *(p)* of 1e-8 per nucleotide per generation (Comeron et al. 2012) and a divergence time *t* of 25 million generations (2.5 million years assuming 10 generations per year) from the common ancestor to each species (Russo et al. 1995). To make the simulations run efficiently we scaled down time and population size by a factor of 1,000 while keeping key parameter products constant to approximate the same evolutionary process (Haller and Messer 2019) (summarized in Table 1).

**Table 1.** Empirical (biological) estimates for evolutionary parameters and scaled estimates used for simulation.

| Parameter | $N_e$ | $\rho$ | $\mu$ | $t$ |
| --- | --- | --- | --- | --- |
| Empirical Estimate | 1e6 | 1e-8 | 2.8e-9 | 2.5e7 |
| Scaled Estimate (simulation) | 1e3 | 1e-5 | 2.8e-6 | 2.5e4 |

$N_e$ : effective population size; $\rho$ : recombination rate; $\mu$ : mutation rate; $t$ : time in unit of generation

We observed 85% of the codons in *bam* encode the same amino acids in both *D. melanogaster* and *D. simulans* reference sequences, likely due to functional constraints (example classifications are show in supplemental table S1). Thus, we used this metric as a baseline in our initial simulations and first randomly sampled 75% of the codons from the identical amino acids in the ancestral sequence to be constrained to the original amino acids. The rest of the identical amino acids (10% of the total amino acids) were assumed to be under completely neutral evolution, while the other 15% unidentical amino acids were subject to selection based on the setup of our models. A nonsynonymous mutation in the conserved codons was always assigned a selection coefficient *s* = −0.1, so that it would undergo strong purifying selection *(N_e_s* = −100 in our simulations). A mutation in the neutral codons was always assigned a selection coefficient *s* = 0.

Each simulation run began with a “neutral burn-in” period of 20,000 simulation generations (=20x scaled *N_e_*) to accumulate genetic variation consistent with an equilibrium state of mutation-drift balance before non-neutral dynamics started. During this period, mutations occurring at the conserved sites were still assigned a selection coefficient of *s* = −0.1 to retain the functionally constrained amino acid positions. At the end of the neutral burn-in period all new variations (fixations and polymorphisms) were retained in the simulation. This quantity of fixations and polymorphisms was checked against the expected number of mutations estimated from population genetic theory based on the given *N_e_, μ*, and the number of (non-conserved) sites in the gene. Selection coefficients of subsequent new mutations were based on a comparison to the original, inferred ancestral sequence.

### Selection regimes

For both the Buffering and Arms Race models we alternated selection on a protein coding gene with and without *Wolbachia* infection. The phases of infection and absence of *Wolbachia* alternated in each model, which simulated the periodic occurrence of *Wolbachia* in natural populations. To keep the simulations simple, we assume that *Wolbachia* infection and loss is instantaneous throughout the entire population and that there are no other effects of *Wolbachia* on the host beyond which we are modeling.

For each selection phase of the simulations, the absolute value of selection coefficient |*s*| for each positively or negatively selected mutation in the 15% of codon sites under selection was fixed for the duration of each simulation. The beneficial mutations were assigned a selection coefficient of *s* > 0 while deleterious mutations had a selection coefficient *s* < 0. To determine the fitness effect of each mutation, we use the amino acid matrix of Miyata et al. (1979).

The Miyata et al. (1979) amino acid matrix captures the primary features of biochemical and physical differences between amino acid pairs. We henceforth refer to the pairwise measures from the Miyata matrix as “Miyata scores (MS)” and use them to determine whether a mutation is neutral (including all synonymous mutations) or under positive or negative selection (nonsynonymous mutations) in each selection scenario. We use a shorthand for Miyata score calculations as follows, with, for example, MS between the current amino acid (AA_cur_) and the mutated amino acid (AA_mut_) represented as MS(AA_cur_, AA_mut_). MS ranges from 0 to 5.13, where 0 indicates identical amino acids, e.g., MS(L,L)=0, and 5.13 indicates the most physicochemically different amino acids, i.e., MS(W,G)=5.13.

Using Miyata scores, we assigned selection coefficients to new mutations in simulated periods of *Wolbachia* infection and absence for each model as outlined in Figure 1. Importantly, the Arms Race model shifts the optimal amino acid sequence to promote amino acid diversification to contrasts it with the Buffering model, which retains an ancestral optimal sequence. The models outlined in Figure 1 are the simplest with regard to assigning selection coefficients and thus are regarded as the “Base” models. For instance, in the Arms Race model, any nonsynonymous mutation would be positively selected during the presence of *Wolbachia.* This allows substitutions to be dominated by amino acids of any type, including those of “small steps” as suggested by Bergman and Eyre-Walker (2019) to be prevalent in *Drosophila.* In order to evaluate the robustness of our simulation results to the choice of amino acid substitution matrix, we also carried out simulations with selection coefficients based on the BLOSUM62 amino acid matrix (Henikoff and Henikoff 1992) which is based on amino acid conservation among protein sequences (relative to random similarity) from diverse taxa with a pairwise identity of no more than 62% (Supplemental Materials and Methods). This measure is differs fundamentally from the Miyata et al. (1970) matrix which considers only the biochemical properties of the amino acids themselves.

**Fig. 1.**
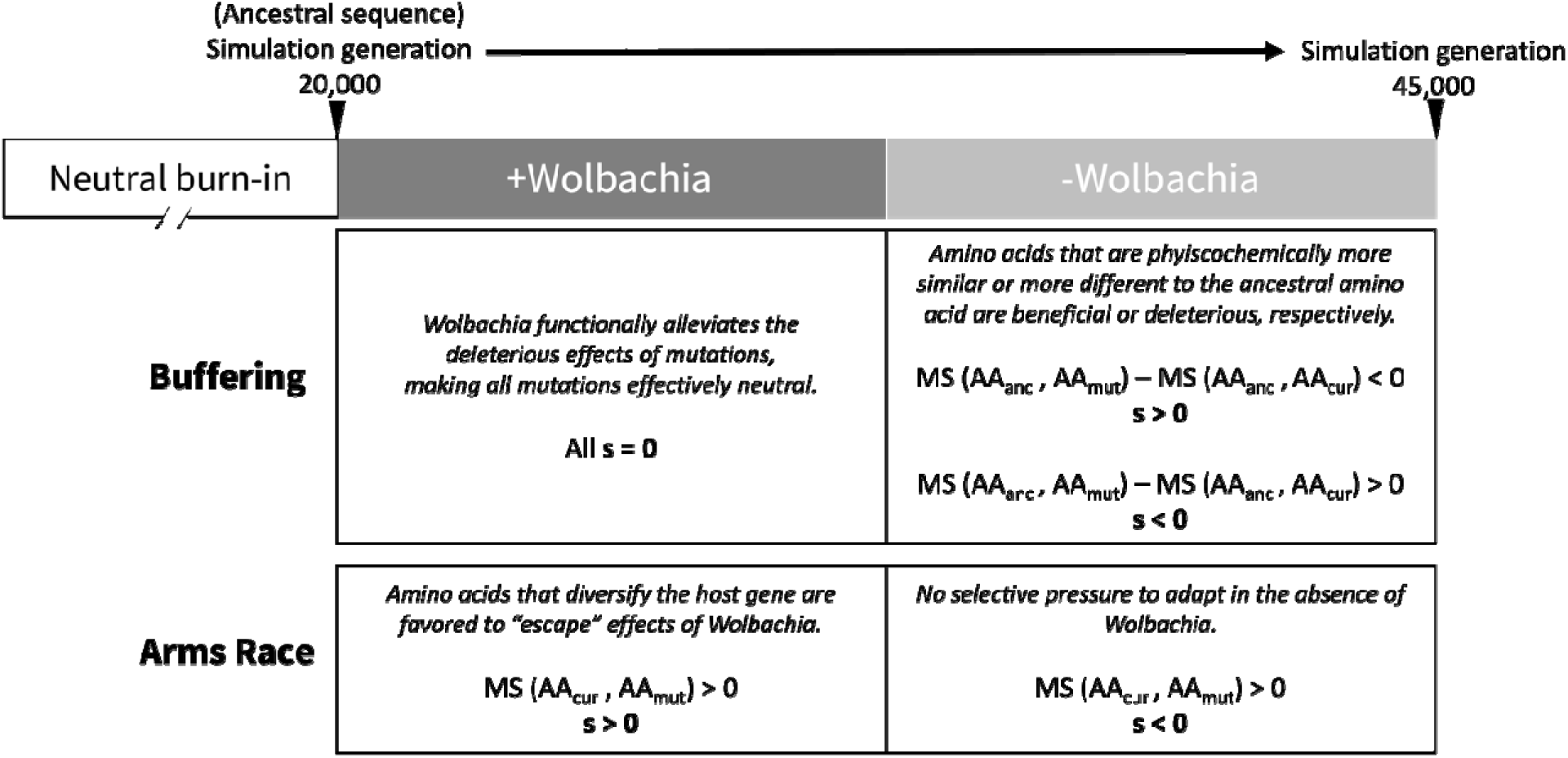
Simulation setup for Buffering and Arms Race Base models. Selection on new nonsynonymous mutations (mutated amino acid, AA_mut_) is determined by their Miyata score (MS) to the appropriate reference amino acid (the current amino acid, AA_cur_, or the ancestral amino acid, AA_anc_).

### Simulation parameters

We focused on investigating the impacts of two key parameters on the evolution of the *Drosophila* species in each of the proposed models: 1) the magnitude of the selection coefficient for both beneficial and deleterious mutations, and 2) the length of alternating *Wolbachia-* infection and *Wolbachia*-absence phases in each model, in which the different selection phases occur. The absolute values of selection coefficients included |*s*| = 0.1, |*s*| = 0.01, and |*s*| = 0.001, resulting in *N_e_*|*s*| = 100, *N_e_*|*s*| = 10, and *N_e_*|*s*| = 1 respectively, where *N_e_*|*s*| = 1 can be considered effectively neutral. The lengths of different selection phases varied from equal periods of 12,500, 6,250, and 3,125 simulation generations (corresponding to 12.5 million, 6.25 million, and 3.125 million generations in unscaled time). For each set of parameter combinations, we ran 50 independent simulations and performed downstream analyses every 3,125 simulation generations after the neutral burn-in period, by comparing the “reference sequence” in SLiM to the common ancestral sequence of *D. melanogaster* and *D. simulans*.

### Analyses of simulated sequences

100 diploid individuals were randomly sampled from the population at the end of each simulation and the numbers of nonsynonymous and synonymous fixations (relative to the inferred ancestral sequence) and polymorphisms present were tabulated. The MK test was used to evaluate departures from an equilibrium neutral model of molecular evolution and was customized from the iMKT package (Murga-Moreno et al. 2019). The Fay et al. (2001) correction (FWW correction) for low frequency polymorphisms was applied, counting only polymorphisms > 5% frequency to avoid including deleterious variation segregating in the populations, and significance was determined by a Fisher’s exact test.

We inferred *α*, which is the proportion of nonsynonymous substitutions driven to fixation by positive selection, in two ways. First, we estimated *α* from sequences sampled at relevant timepoints in the simulation following Smith and Eyre-Walker (2002) 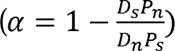, where D_s_ and D_n_ are the number of fixed synonymous and nonsynonymous substitutions and P_s_ and P_n_ are the number of polymorphisms in the sample that are synonymous and nonsynonymous. We refer to this sample derived statistic as the “estimated *α*” throughout this paper, and this is the only type of estimate of *α* that can be obtained by experimentalists for samples of sequences from populations.

For comparison, we also calculated the “true *α”* in the simulations by tracking the actual proportion of all nonsynonymous substitutions in *bam* that were driven to fixation by positive selection in the simulation. Thus true *α* = (number of nonsynonymous mutations that fix and have s > 0)/(all nonsynonymous substitutions). Since the selection coefficient of a mutation could have changed as *Wolbachia* was gained and lost from the population, any mutation that once had a selection coefficient *s* > 0 and was eventually fixed in the population was regarded as being driven to fixation by positive selection. As with the estimated *α*, the true *α* is calculated for each simulation from the observed substitutions relative to the ancestral sequence.

Lastly, the average Miyata score calculated for each amino acid change between the simulated, evolved sequence and the ancestral sequence was used as an assessment of physicochemical similarity between the two sequences.

## Results

### Buffering Model

We first simulated the Buffering Base model based on the *Drosophila bam* gene, together with the cyclic pattern of *Wolbachia* infection and loss in the *Drosophila* population. We had predicted that slightly deleterious nonsynonymous mutations would be functionally buffered by the presence of *Wolbachia* such that these mutations become effectively neutral and could drift to fixation. However, when *Wolbachia* is lost, new mutations that brought the *bam* protein closer to the ancestral functional state would be positively selected, while any mutation that pushed the *bam* protein further away from the ancestry would be selected against.

Simulations demonstrated this prediction to be true: some nonsynonymous substitutions were driven to fixation by positive selection as measured by “true *α*” in the first *Wolbachia-* absence phase (fig 2, row 1). This pattern was most evident with the strongest selection: the true *α* increased and stayed constant in the phases where positive selection was expected, with only a marginal decrease during the phases of neutral evolution. Weaker selection led to a smaller increase in the true *α*. Notably, longer *Wolbachia* infection periods resulted in larger true *α*’s, presumably due to the longer time to accumulate buffered deleterious fixations by drift in the presence of *Wolbachia*.

**Fig. 2.**
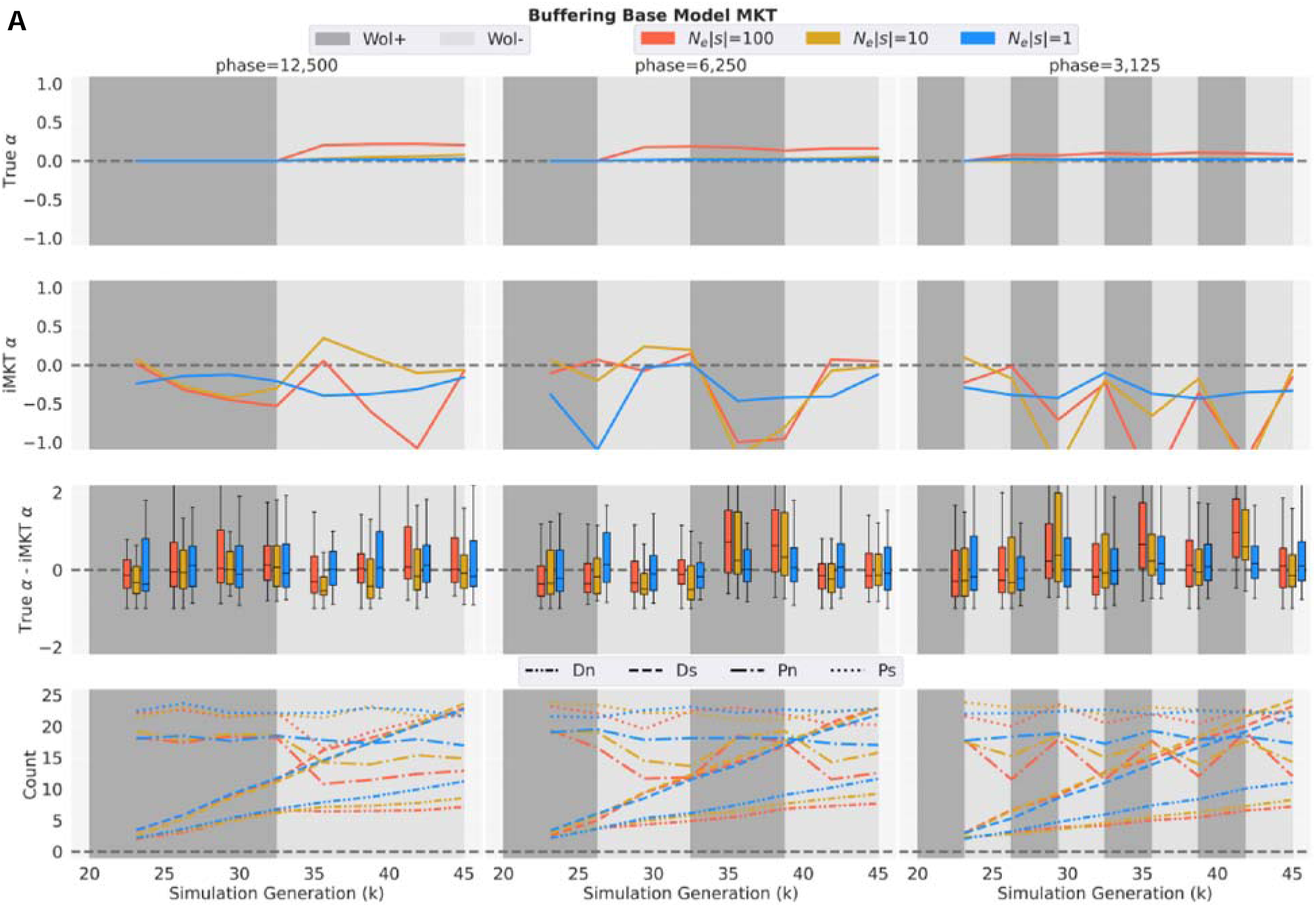
MK test results of simulations for Buffering Base model. Each panel shows MK test (MKT) analyses with different selection coefficients of *N_e_|s|*=100, *N_e_|s|*=10, and *N_e_|s|*=1 graphed across alternating phases (phase length=12,500, 6,250, and 3,125 simulation generations) of *Wolbachia* infection (Wol+, dark grey) and *Wolbachia* absence (Wol-, light grey) post-burn-in period. In each panel, row 1: the average true *α* in the simulations; row 2: the average estimated *α* (iMKT *α*) in the simulations (FWW correction, SNPs frequency > 5% only); row 3: the distributions of differences between the true and estimated *α* every 3,125 simulation generations; row 4: The average of each MK test component (D_n_, D_s_, P_n_, P_s_).

Despite the pattern for true *α*, estimated *α*’s from the observed numbers of nonsynonymous and synonymous fixations and polymorphisms in the samples of sequences from the simulations were largely negative across the whole simulation in the Buffering Base model regardless of the selection coefficients (fig. 2, row 2). Negative estimated a’s indicate a violation of MK test assumptions and are, here, a result of high P_n_ and low D_n_. In the initial *Wolbachia*-infection phase, nonsynonymous polymorphisms were negatively selected in the constrained codons and neutrally buffered by *Wolbachia* in the codons under selection, with few such mutations in the latter category going to fixation (fig 2, row 4). Following this “buffering” period, a subset of nonsynonymous mutations was selected for. However, the number of nonsynonymous mutations that could be positively selected in the Buffering Base model was limited, leading to a smaller D_n_ and thus a smaller (possibly < 0) estimated *α*, even when positive selection was present as evidenced by the true *α*. Overall, estimated *α* decreased during neutral phases *(Wolbachia* present and thus a phase of genetic drift) but increased in phases with selection *(Wolbachia* absent and thus selection to return *bam* to a more functional state).

The boxplots of differences between the true and estimated a’s illustrated that estimated a’s systematically underestimate the true *α*’s. This is due to the presence of deleterious polymorphisms (Fay et al. 2001, Eyre-Walker and Keightley 2009, Messer and Petrov 2013), which is what we observe here with the boxplots distributed above 0 (especially with *A^r^_e_*|*s*|>1 in the later *Wolbachia*-infection phases). For the four MK test parameters, the final magnitude of D_n_, D_s_, P_n_, and P_s_ observed at the end of the simulations were slightly impacted by the length of *Wolbachia* infection and absence periods, but only P_n_ showed dramatic periodic fluctuations due to the cyclic infection and absence periods (fig. 2, row 4).

Additionally, we looked at the distributions of p-values from the MK test (FWW correction, SNPs frequency > 5%) and the correlation between them and the estimated a’s at the end of the simulation. We find that even under the strongest selection in our simulations, the MK test could hardly detect any statistically significant signals of positive selection in the Buffering Base model, likely due in part to the modest length of the *bam* gene (fig 4A). Overall, smaller p-values were associated with larger estimated a’s, and all the significant p-values (p<0.05) were associated with estimated a’s close to 1.0 across all selection coefficients (data not shown).

All together, these results demonstrate that a modest number of amino acid fixations can occur due to selection for an optimal ancestral allele after a period of buffered mutations accumulate. The estimated *α* did not, however, reliably identify departures from neutrality in the direction of positive selection in the Buffering Base model.

To further explore the Buffering model and its potential to generate signals of positive selection, we also ran additional simulations with different Miyata score cutoffs informing the selection schemes (Supplemental Materials and Methods). Here, we introduced neutral and deleterious ranges to allow for a more nuanced selection scheme and termed the new model as the Buffering Complex model (fig. S1). The Buffering Complex models had lower true *α*’s compared to Base models (fig. S2, row 1), with a barely perceptible increase in true *α* when *Wolbachia* is lost from the population, even in the strongest selection scenario of *N_e_\s\ =* 100. The patterns of estimated a’s and boxplots for the difference between the true and estimated a’s were similar between Complex and Base models (fig S2, row 2&3). D_n_ had a slight increase in the Buffering Complex model compared with the Buffering Base model, potentially due to the introduction of the neutral region leading to a small number of additional nonsynonymous fixations by genetic drift alone (fig S2, row 4). The MK test still could detect only a minimal number of statistically significant signals of positive selection in the Complex model (fig. S4A), just like the case for the Base model.

These results suggest that restricting the degrees to which we positively select an amino acid based on their physicochemical change makes it harder to generate signatures of positive selection for *bam* in the Buffering models, especially with a limited infection period. The signatures of positive selection required both fixations of deleterious mutations during the infection and a significant amount of amino acids to be positively selected due to their ability to revert mutated proteins back to their optimal functions.

### Arms Race Model

We compare our Buffering model with that of a traditional arms race model between two species, which we implemented with positive selection acting on nonsynonymous mutations in the presence of *Wolbachia* and purifying selection imposed on nonsynonymous mutations in the absence of *Wolbachia.* We find that the patterns of true *α*’s were clearly indicative of positive selection in the Arms Race Base model phase with *Wolbachia* (fig 3, row 1). The elevated true *α*’s persisted, though with a slow decline during the subsequent *Wolbachia-free* phase. Positive selection in the *Wolbachia* infection phases increased *α* or kept it as a constant, while purifying selection in the *Wolbachia* absence phases decreased *α* marginally.

**Fig. 3.**
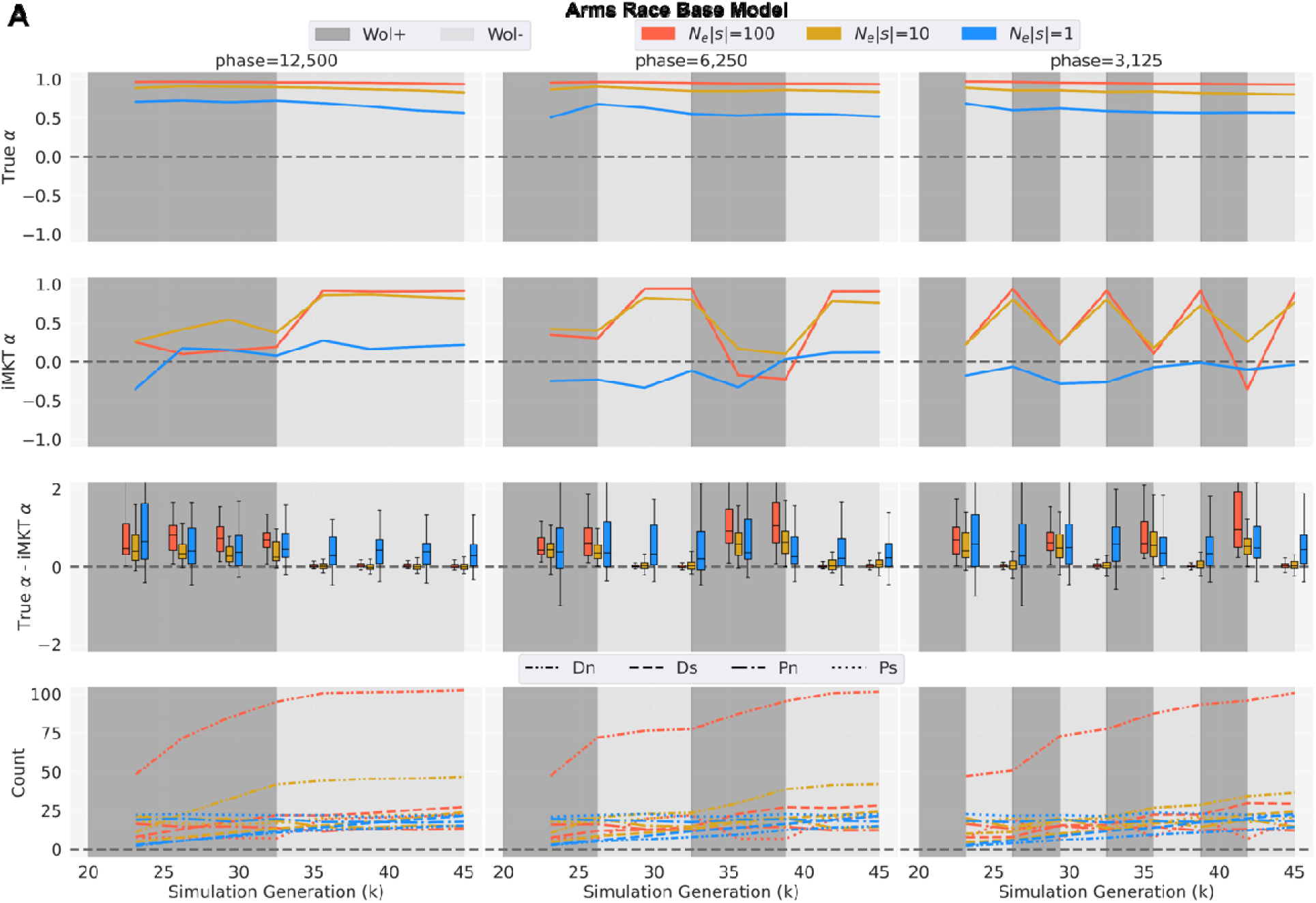
MK test results of simulations for Arms race Base model. Each panel shows MKT analyses with different selection coefficients of *N_e_|s|*=100, *N_e_|s|*=10, and *N_e_|s|*=1 graphed across alternating phases (phase length=12,500, 6,250, and 3,125 simulation generations) of *Wolbach**ia*** infection (Wol+, dark grey) and *Wolbachia* absence (Wol-, light grey) post-burn-in period. In each panel, row 1: the average true in the simulations; row 2: the average estimated (iMKT) in the simulations (FWW correction, SNPs frequency > 5% only); row 3: the distributions of differences between the true and estimated every 3,125 simulation generations; row 4: The average of each MK test component (D_n_, D_s_, P_n_, P_s_).

The averages of estimated a’s were almost all positive for selection coefficients with *N_e_*|*s*|>1 and showed clear periodic changes as *Wolbachia* comes in and out of the population across all three phase lengths (fig. 3, row 2). Surprisingly, the magnitude of the estimated a’s increased in the phase without the imposed positive selection. This unexpected increase is explained by the change of nonsynonymous polymorphisms in the population. In the *Wolbachia-* infection phase, both D_n_ and P_n_ accumulated due to the positive selection of nonsynonymous mutations, as expected; however, after the sudden change to the *Wolbachia-absence* phase, D_n_ was largely unchanged while P_n_ experienced a sudden decrease as segregating and new nonsynonymous mutations were all selected against (fig 3, row 4). Given the equation for estimating a from a sample of sequences 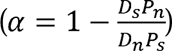, the estimated *α* therefore increased in the phase with the implemented purifying selection that followed the positive selection. For the effectively neutral case of *N_e_\s\* = 1, the estimated *α*’s in the Arms Race Base model fluctuated around 0.

For the Arms Race Base model with effectively neutral evolution (*N_e_*|*s*|=1), estimated a’s usually underestimated the true *α*’s (fig 3, row 3). However, under stronger selection (*N_e_*|*s*|=10 or 100), estimated a’s underestimated the true *α*’s only during the *Wolbachia*-infection phase; there was good accuracy in estimated a’s estimation when *Wolbachia* was lost, which reflected the delay in detecting selection based on changing P_n_ as previously explained. The pattern of the true *α*’s and the estimated a’s was not dramatically influenced by the magnitude of the selection coefficient (*N_e_*|*s*|=10 or 100) or the varying lengths of the infection/absence periods that we examined.

We found that a statistically significant rejection of neutrality in the direction of positive selection was more likely to be detected with the MK test in the Arms Race Base model than in the Buffering Base model, since the values of the key MK test parameter D_n_ are generally much larger in both *Wolbachia*-infection and *Wolbachia*-absence phases in the Arms Race model. This increased magnitude of D_n_ provided more statistical power in Fisher’s exact test (fig. 4B). Overall, smaller p-values were always associated with larger estimated a’s, and all the significant p-values (p<0.05) were associated with estimated a’s close to 1.0 across all selection coefficients (data not shown).

**Fig. 4.**
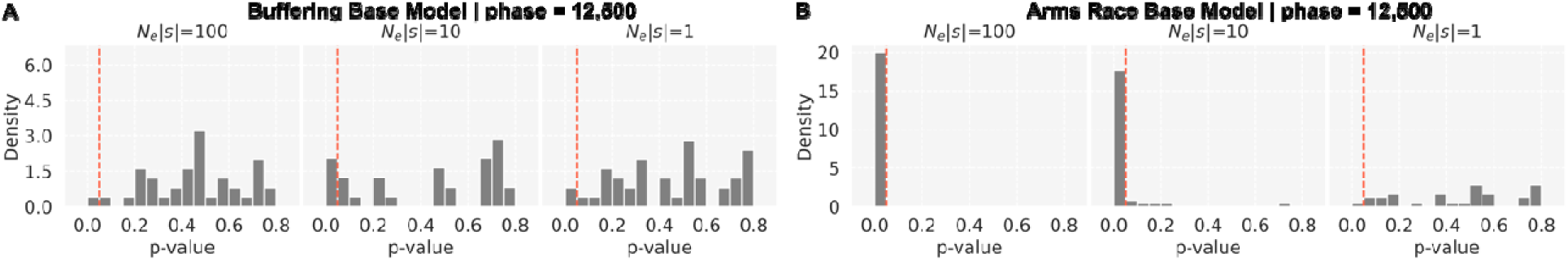
Distributions of MK test p-values for Base models. MK test p-values (FWW correcti**on**, SNPs frequency > 5% only) for simulation runs with *Wolbachia* phase=12,500 simulation generations and *N_e_|s|*=100, 10, and 1 for the simulated models at 45,000 simulation generation. The vertical red line denotes p-value=0.05. Note that distributions are normalized to have an ar**ea** of 1 under the histograms. (A) Buffering Base model; (B) Arms Race Base model.

As with the Buffering model, we implemented the Arms Race model with different Miyata score selection-based cutoffs and termed it as the Arms Race Complex model (Supplemental Materials and Methods; fig. S1). The MK test results for the Arms Race Complex model closely resembled those for the Base model. Since we narrowed down the Miyata score range for the positively selected nonsynonymous mutations by introducing neutral and deleterious ranges, we observed lower true *α*’s compared to the Base model (fig S3, row 1). The patterns of estimated a’s and boxplots for the difference between the true and estimated a’s were similar between Complex and Base models (fig S3, row 2&3). The total number of D_n_ did not reach the same magnitude at the end of simulation for the Arms Race Complex model as it did in Arms Race Base model across different phase lengths and selection coefficients (fig S3, row 4). However, the MK test still detects statistical signals of positive selection under the first two selection coefficients in the Complex model (fig. S4B).

### Distributions of Miyata scores under Buffering and Arms Race models

We have observed that Buffering models can result in fixation of amino acids due to positive selection based on true *α*’s, but the estimated a signals were harder to detect compared to the cases in the Arms Race models. We expect amino acid substitutions to be more diversified in the Arms Race model than in the Buffering model in both the Base and Complex cases, as the former is based on the premise of a sequence evolving *away* from the ancestral sequence and the latter is based on the premise of a sequence evolving *toward* the ancestral sequence. To assess this, we calculated Miyata scores between each amino acid substitution and its ancestral amino acid of the post-burn-in SLiM reference sequence to represent their physiochemical differences.

For both the Base and Complex cases, the distributions of Miyata scores per amino acid from the Arms Race and Buffering models were most distinguishable from each other in the strongest selection scenario at *N_e_\s\ =* 100. Here, the interquartile ranges of Miyata score distributions of the two models were completely separated at the end of simulations (fig 5, row 1; fig. S5, row 1), but they were basically indistinguishable from each other throughout the simulations when the selection is the weakest at *N_e_\s\ =* 1 (fig. 5, row 3; fig. S5, row 3).

**Fig. 5.**
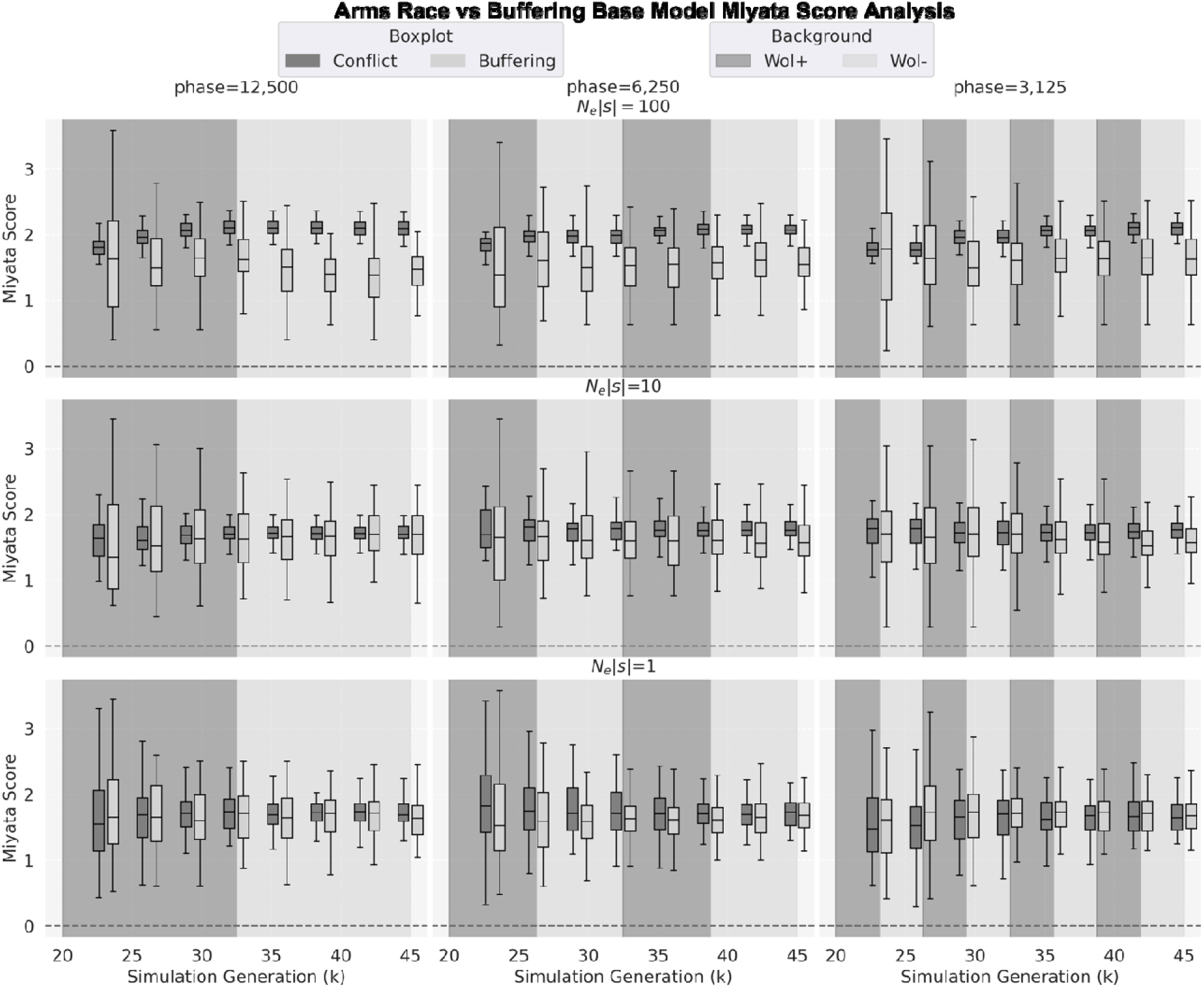
The distribution of Miyata scores per amino acid substitution for Base models. Miyata scores per amino acid substitution across multiple runs for substitutions between the consensus sequence at the end of a given simulation generation and the ancestral sequence. Data is shown for both the Arms Race model (dark grey) and Buffering model (light grey) at every 3,125 simulated generations post burn-in for different phase lengths.

For *N_e_\s\ =* 10 and *N_e_\s\ =* 100, the distributions of Miyata scores overlap more in the Base case than in the Complex case for both Buffering and Arms Race models (fig 5, row 2 fig. S5, row2) because the positively selected mutations had a higher concentration of Miyata scores between 1 and 3 in the Complex case, which made the differences between Miyata scores more prominent. Different infection/absence phase lengths did not have a large impact on the average Miyata scores across the simulations.

### BLOSUM62-based simulations

We complemented the above simulations with another simulation setup that assigned selection coefficients based on the BLOSUM62 amino acid matrix (Henikoff and Henikoff 1992) (fig S6). MK test results were largely consistent with the results from the Miyata-score-based Buffering and Arms Race Base simulations, again showing that the positive true *α* values that are produced in the Buffering model are not well captured by the estimated *α* (fig S7). The associated p-values were likewise similar to the Miyata-based simulations, but with one striking difference: The *N_e_\s\* =100 BLOSUM62-based Arms Race model had much fewer statistically significant results (fig. S8). This is due to the high number of low frequency polymorphisms that accumulate in the BLOSUM62-based simulations that are removed with the FWW correction (fig S9).

### Comparison with the empirical data

To evaluate which model in our analysis better captures *bam’s* observed patterns of sequence evolution within and between natural populations of *Drosophila*, we first performed the MK test on a population sample (n=89) of *D. melanogaster* (Lack, et al. 2015), using divergence to the predicted common ancestral sequence with *D. simulans* as the outgroup and a randomly sampled sequence as the reference sequence used in the estimated *α*. Analysis of these data reject neutrality in the direction of positive selection using the MK test with a p = 0.00015 and estimated *α* of 0.91 (FWW correction, SNPs > 5% only). We used the number of nonsynonymous substitutions per nonsynonymous site (dN) calculated from MK test results as the summary statistic to tune selection parameters of the two simulation models with only one *Wolbachia* infection-loss cycle.

While examining polymorphism levels would seem important to distinguish between Buffering and Arms Race models, these levels are very sensitive to the length of *Wolbachia* infection and absence as we have modeled it, for example, due to the strong purifying selection occurring in the Arms Race model when *Wolbachia* is lost. The problematic effect of this timing choice on P_n_ and P_s_ can be seen in Figures 2 and 3. As D_n_ is less sensitive to the sampling time points and represents the number of amino acid changes in *bam*, we chose to only use this parameter to evaluate how well our models fit to empirical data.

Applying the MK test to our empirical *D. melanogaster* population data, we found that D_n_ = 34 and dN = 0.033. We initially found that Arms Race models always predicted a much higher D_n_ than the empirical observation, while Buffering models often exhibited a much lower D_n_. Such results showed the initially assumed ratio of codons under selection (RS=15%) and ratio of codons under constraints (RC=75%) could not reproduce similar results for the evolution of amino acids in either model. Thus, we chose to tune these two ratios of selected and constrained codons (RS and RC) under different strengths of positive selection (*N_e_|s*| = 100, 10, 1) and explore under which parameter settings could we fit the empirical dN. We maintained the mutation rate, population size, and recombination rate when fitting the models to the empirical data because they were based on empirical estimates. When we achieved a matching dN, we then compared the Miyata scores per amino acid change in the observed data and our simulation results to see whether an Arms Race or a Buffering model is more similar to our empirical observations.

For each selection coefficient *s*, we ran simulations using RS and RC both sampled from a uniformly distributed grid of nine points ranging from 0 to 0.8, since the maximum proportion of conserved codons is 0.85, and assessed the resulting dN. For the Arms Race models, dN was consistently more than two-fold overestimated for any proportion of selected sites greater than 0 (e.g., RS > 0, data not shown). Therefore, we refined the Arms Race model grid search for RS to a uniform grid of 6 points from [0, 0.1], while keeping the full grid range for RC. For the Buffering models, we kept the full range of the RS grid as we did find parameters that fit the observed dN. For each pair of parameters, we ran 50 simulations and calculated the mean of dN (dN) across the runs. We then compared the difference between the empirical dN and dN. The best pair of RS and RC was the one that led to the smallest difference between dN and the empirical dN under each selection coefficient *s*.

For the models with selection coefficient *N_e_\s\* = 1, all combinations of the two ratios reproduced similar results consistent with effective neutrality (fig. S10). For moderate or strong selection, the best-fit parameters are shown in Table 2.

**Table 2.**
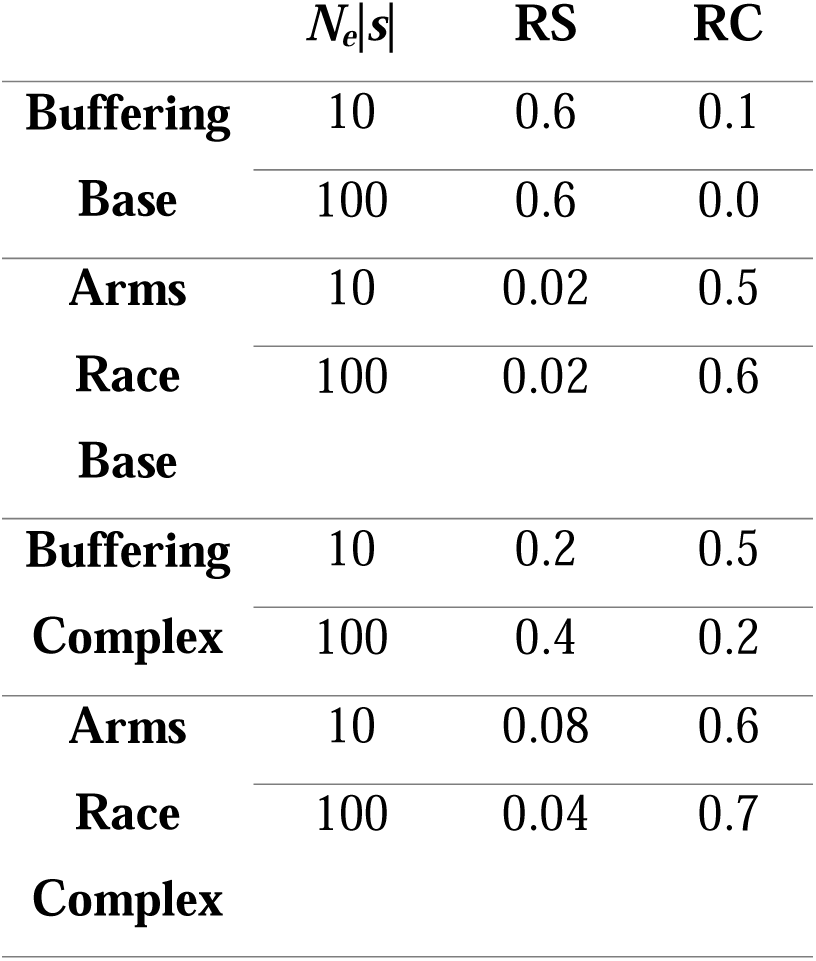
Best fit RS and RC parameters for Ne|s| =10 and 100 for Buffering and Arms race models.

### Analyses of Buffering and Arms Race Models Best Fitting the Empirical Data

To evaluate how well the Buffering and Arms Race models implemented with the best-fit pairs of RS and RC recapitulate the empirical data for *D. melanogaster*, we performed the same MK test and Miyata score analysis for the resulting simulations. Positive true *α*’s were observed in the Buffering Base, Arms Race Base, and Arms Race Complex models across different phase lengths, indicating that positive selection was present under these scenarios. However, the MK test could only identify positive selection by the estimated a and statistically significant p-values in the two Arms Race models with strong selection at *N_e_\s\* = 100. Moderate selection at *N_e_\s\* = 10 in the Arms Race models or any levels of selection in the two Buffering models was not detected by MK test p-values or estimated a (fig. 6; fig. S11).

**Fig. 6.**
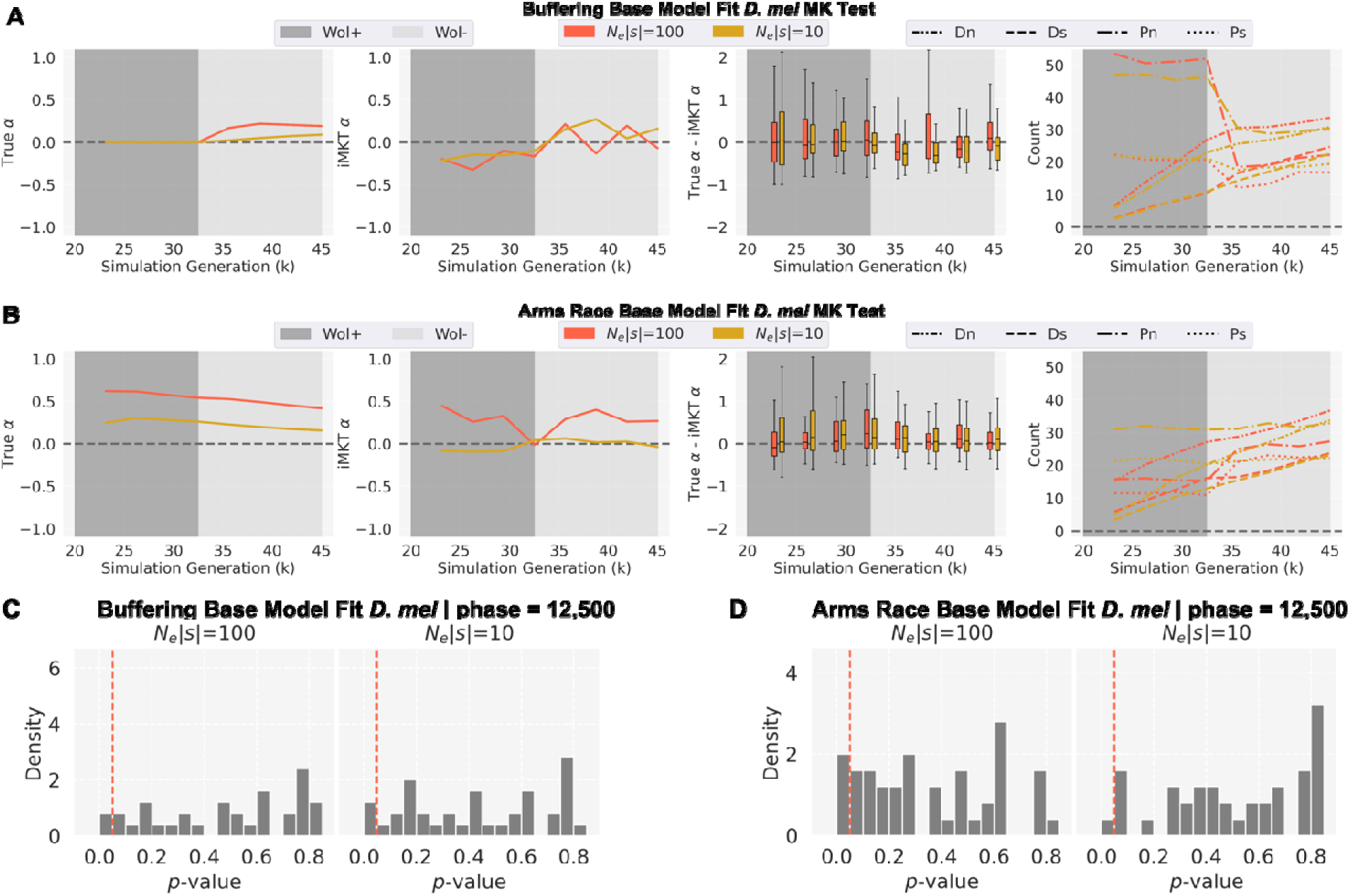
MK test results of simulations for best-fit Base models for *D. melanogaster*. (A-B) analysis of each model with different selection coefficients of *N_e_|s|*=100 and *N_e_|s|*=10 graphed at phase length=12,500 simulation generations of *Wolbachia* infection (Wol+, dark grey) and *Wolbachia* absence (Wol-, light grey) post-burn-in period. In each panel, column 1: the average true in the simulations; column 2: the average estimated (iMKT) in the simulations (FW**W** correction, SNPs frequency > 5% only); column 3: the distributions of differences between the true and estimated every 3,125 simulation generations; column 4: The average of each MK te**st** component (D_n_, D_s_, P_n_, P_s_). (C-D) Distributions of MK test p-values. MK teset p-values (FWW correction, SNPs frequency > 5% only) for simulation runs with *Wolbachia* phase=12,500 simulation generations and *N_e_|s|*=100 and 10 for the simulated models at 45,000 simulation generation. The vertical red line denotes p-value=0.05. Note that distributions are normalized to have an area of 1 under the histograms.

In addition, we calculated the empirical per-site Miyata scores between the current *D. melanogaster* sequences and their predicted common ancestral sequence shared with *D. simulans* (supplemental file S2) and compared it with the distributions of per-site Miyata scores simulated from the best-fitted RS and RC at different timepoints. The end of the simulations at 45,000 scaled generations represents the actual divergence time between the ancestral sequence and the extant *D. melanogaster* and *D. simulans* species. At this time point, the interquartile ranges of Miyata scores of the Buffering and Arms Race models have separated from each other, with fully non-overlapping interquartile distributions in the Complex models. In all models, the per-site Miyata score of *D. melanogaster* are located closer to the center of the distributions from the Buffering models than to the center of distributions from the Arms Race models (fig. 7; fig. S12).

**Fig. 7.**
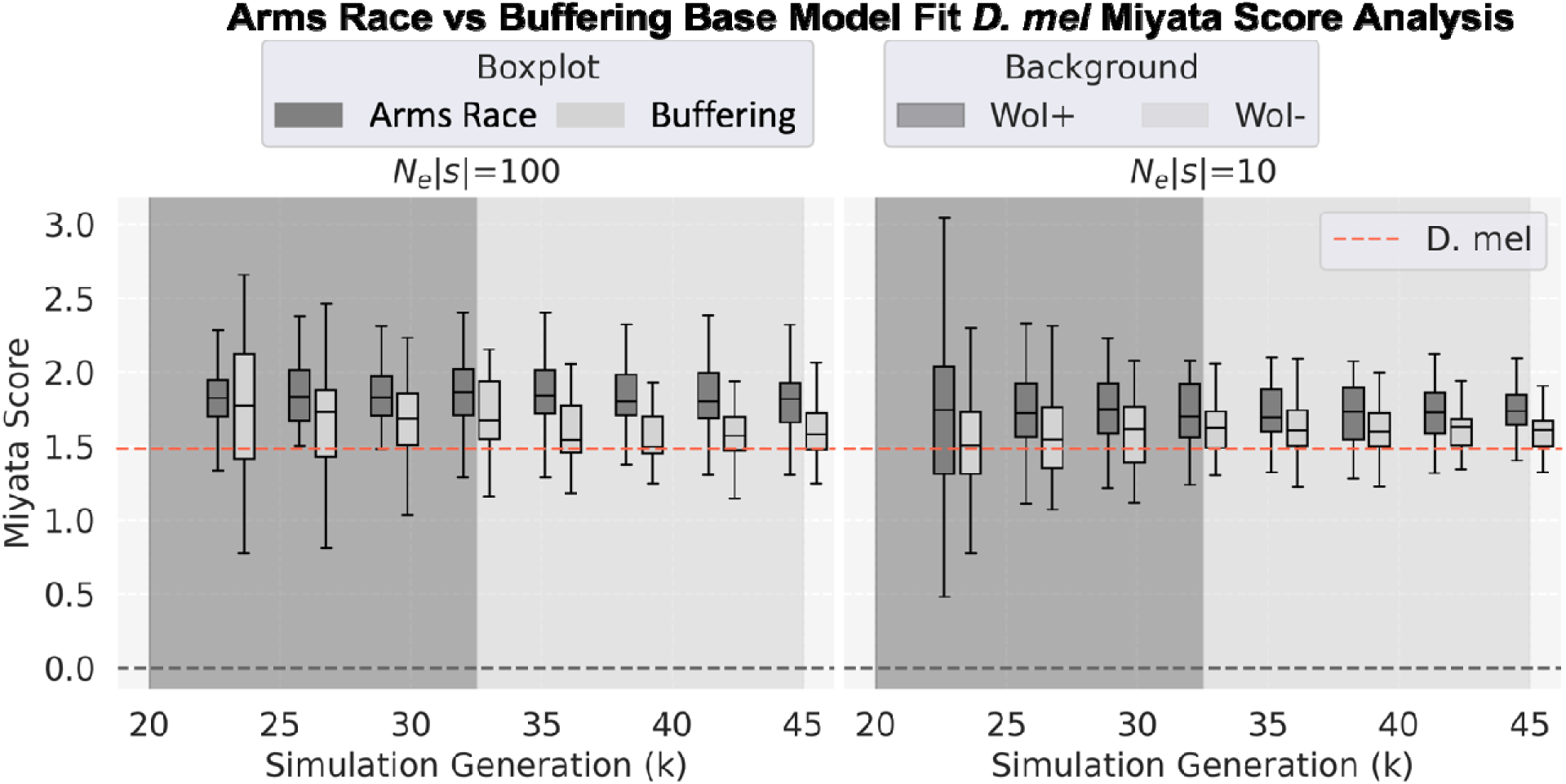
Distributions of Miyata score for the *D. melanogaster* samples from the simulations with best-fit parameters for Base models. The boxplots are the distributions of Miyata scores for each model at every 3,125 generation. The distributions are compared with the observed summary statistics of *D. melanogaster* empirical data (red horizontal line).

In summary, the best-fit Arms Race models with strong selection reproduced the most significant MK test p-values and high estimates of a like those observed in the *D. melanogaster* sample, but the Miyata-score analysis indicated the Buffering models as a better fit for the evolution of the amino acids’ biochemical properties. It is important to note that while the average estimated a is close to zero for all Buffering models, the lower whiskers on the box plots in fig 6A and 6C show that high MK test estimated a’s can, although infrequently, occur under the Buffering models as well.

## Discussion

We have evaluated the molecular evolutionary impact of a Buffering model based on the observations that *Wolbachia* protects the functions of *bam* and *Sxl* from the effects of deleterious mutations, thereby allowing these mutations to accumulate during the *Wolbachia* infection phase by drift (equivalent to the relaxation of functional constraints for amino acid mutations). When *Wolbachia* is lost, the constraints are reimposed and amino acids similar to *bam* and *Sxl*’s ancestral state are positively selected for, potentially leading to a signature of positive selection in a sample of the resulting DNA sequences. We compared predictions of this model to those from an Arms Race model, which is an implementation of dynamics that may describe the interactions between, for example, a host and parasite.

We used simulations to study the evolutionary process involved in each model and found that both models can generate positively selected amino acid fixations, as measured by the actual number of such fixations tracked in our simulations (the “true *α*”). A positive true *α* in the Buffering models reveals that *Wolbachia* need not function as a reproductive parasite in arms race with a gene like *bam* to drive positive selection in the host gene.

Importantly, in all simulations the estimated *α* generally underestimated the true *α*. This underestimation had a minimal effect on our interpretation of evolution in the Arms Race models, as the true *α* was very large in all simulations outside of those with the weakest selection. On the other hand, with a maximum true *α* of ∼0.25 in the Buffering models’ simulations, an underestimation led to a weak or absent signal of positive selection detectable by the MK test, which could further be confounded by statistical noise. Such findings highlight some limitations of the MK test noted by others (Akashi 1999, Fay et al. 2001, Eyre-Walker and Keightley 2009, Zhai et al. 2009, Messer and Petrov 2013). Even with these limitations, the MK test could still infer high a’s in some simulation runs under the Buffering models, representing detection of positive selection.

The Buffering models require the initial fixation by genetic drift of *Wolbachia*-buffered deleterious nonsynonymous mutations for there to be resulting positive selection during a subsequent phase without *Wolbachia.* This effect is seen across the three different infection lengths that we simulated in the Buffering Base model. Longer *Wolbachia* infection phases increase the chance of detecting positive selection in a subsequent *Wolbachia* absence phase, though never to the level resulting from Arms Race models. The average length of *Wolbachia* infection time is unknown for *Drosophila*, but two independent studies that found the *w*Mel *Wolbachia* variant to have been in *D. melanogaster* for at least 79,000 and 80,000 *Drosophila* generations (Richardson et al. 2012; Choi and Aquadro, 2014). These time periods are shorter than what we have simulated, but there is evidence to suggest turnover of *Wolbachia* variants that could act as a longer standing infection period than currently documented (Riegler et al. 2005, Kriesner et al. 2013). Thus, *Wolbachia* infection of the length we have simulated, and with it a potential for subsequent positive selection, is not out of question.

To better evaluate the fit of the observed data from *D. melanogaster* to the predictions of the Buffering and Arms Race models, we tuned the simulation selection parameters of both models to fit the observed nonsynonymous sequence divergence per nonsynonymous site (dN) between *D. melanogaster* and the inferred common ancestor with *D. simulans.* Only the tuned Arms Race model recapitulated the statistically significant positive estimated a’s that we observed for the *D. melanogaster* population. As in the general Buffering results discussed above, the tuned Buffering model resulted in evidence of positive selection as indicated by a true *α* under certain conditions, but we could rarely detect it with the MK test in *bam* with statistical significance. For the Miyata score analysis, we found that the Buffering models better fit our empirical data, as the Arms Race models predicts greater amino acid diversity than we observe. We expect that a restriction of the positively selected amino acids in the Arms Race model to only those physicochemically similar to the current states would bring the Miyata score analysis in line with our observed amino acid diversity without compromising the high estimated a that is generated. Thus, combining these results, we suggest that the Buffering model is a possible, but unlikely, explanation behind the observed evolution in the *D. melanogaster bam* gene. This is particularly the case as a p-value less than 0.05 for the empirical MK test result is the typical criteria used by experimentalists to infer a departure from an equilibrium neutral model. Thus, with the current assumptions of our models, the Arms Race is the better explanation for the signature of selection that we observe at *bam.* Of note is that a change in function that favors diversification of the protein-coding gene would give similar results because, like the Arms Race model, selection to refine a new function would likely favor positive selection for physicochemically different amino acids. A recent analysis of CRISPR/Cas-9 generated nulls in five *Drosophila* species raises this possibility for *bam* (Bubnell et al. 2022). Whether the observed changes in function are associated with arms race with *Wolbachia* remains an open question as the two are not mutually exclusive.

We note that the best fit results for all models come with parameterizations that include a considerable proportion of neutral sites. This suggests that our model is missing important subtleties behind the evolution of *bam*. For instance, we have only used fixed selection coefficients and strict Miyata score cutoffs throughout our simulations to model the selection coefficients for both beneficial and deleterious mutations, when they could be drawn from some distribution. It is also possible that a mixture of Buffering and Arms Race models may be operating, with each driving evolution at a subset of sites.

With regard to resolving the evolutionary interactions between *bam* and *Wolbachia* in *Drosophila*, it will now be important to explore other experimental evidence with respect to potential arms race, change in function, or buffering effects due to relaxed selective constraints. For example, we can further evaluate the potential contributions of an arms race by testing for positive selection in *Wolbachia* genes. In an Arms Race model, *Wolbachia* would co-evolve with *bam* to continue its impact on *Drosophila* fertility. There is already some evidence of positive selection across different *Wolbachia* strains of arthropods and nematodes (Baldo et al. 2002; Baldo et al. 2010) but a much more thorough analysis of closely related *Wolbachia* strains infecting *D. melanogaster* and its close relatives is needed.

The applicability of the Buffering model extends beyond the *bam* gene in *Drosophila.* Other cases of *Wolbachia* interacting with the *Drosophila* germline include the rescue of *Sxl* hypomorphs (Starr and Cline 2002) and one *mei-P26* hypomorph (unpublished). *Wolbachia* also increases the fecundity of *D. mauritiana* (Fast et al. 2011). These examples suggest *Wolbachia* could function in a “buffering” manner to increase the fitness of its host across a variety of situations, especially given the high prevalence of *Wolbachia* across different species (Hilgenboecker et al. 2008; Zug et al. 2012; Weinert et al. 2015). The Buffering model may also be pertinent to other facultative symbiotic relationships wherein one organism acts as a defensive symbiont by providing protection from a natural enemy for the other species (reviewed in Clay 2014).

Species with smaller population sizes may present better opportunities for detection of positive selection from the Buffering model. Drift during the “buffering” phase is what allows the buffered deleterious nonsynonymous mutations to fix and the time to fixation of neutral mutations is approximately *4N_e_* generations (Kimura and Ohta 1969). Thus, a smaller *N_e_* will allow for more chances of deleterious mutations fixing and subsequently being under selection to return to an ancestral state. The effectiveness of selection is also reduced, however, by a smaller *N_e_.* Genes of a greater length may also provide more statistical power to detect positive selection.

In conclusion, the Buffering model is based on population genetic theory where there is episodic variation between positive and relaxed selection on a protein coding gene. Our results motivate consideration of buffering-like evolutionary processes when population genetic evidence is found for departures from selective neutrality consistent with positive selection. For example, the framework that could apply to populations that experience cycles of higher mutational loads, followed by positive selection. This could be observed in temperate species affected by ice ages, where a drop in population size allows the fixation of some deleterious alleles that are subsequently purged during population expansion after the ice age retreats.

The Buffering model sits well alongside previously proposed ideas of conditional neutrality and antagonistic pleiotropy that offer dynamics of local adaptation resulting from a genotype x environment interaction. With conditional neutrality, an allele varies between neutral and high fitness across two different environments. With antagonistic pleiotropy, two alleles reciprocally vary between high and low fitness across two different environments. The Buffering model offers an intermediary scenario, wherein one allele varies between neutral and low fitness across two different environments and the other varies between neutral and high fitness across two different environments. As we have shown that a buffering dynamic can produce selectively fixed amino acids, genomic signatures and statistical tests that distinguish buffering-driven adaptation from other forms of adaptation, including arms race dynamics, should be further examined.

## Supplemental Materials

Supplemental materials are available at __________.

## Data Availability

No new sequencing data were generated in this work; the employed data sets are listed throughout the text. Sequence alignments used are available as supplementary file S2, Supplementary material online. SLiM3 and python code for analyses used in this study are available online at github.com/runxi-shen/Modeling-Evolution-at-bam.

## Notes

### Competing Interest Statement

The authors have declared no competing interest.

### Summary of Updates

Additional simulations and analyses have been performed and the presentation of the story has been reframed and streamlined for ease of reading.

