## Supplemental Materials and Methods for "A model of functionally buffered deleterious mutations can lead to signatures of positive selection"

**BLOSUM62-based selection schematic**

To test the robustness of our models, we perform additional simulations using the BLOSUM62 amino acid matrix as the metric informing whether a mutation is neutral or under positive or negative selection (fig S6). Unlike the Miyata matrix, the BLOSUM62 matrix scores have no regard to the physiochemical nature of the amino acids. The scores represent the odds of finding an amino acid pair aligned in the designated database of homologous proteins (Henikoff and Henikoff 1992). Nomenclature will follow like that for the Miyata matrix, where pairwise measures from the BLOSUM62 matrix are referred to as “BLOSUM62 scores (BS).”

**Complex models**

We test “Complex” models that limit the mutations under positive selection under the preposition that mutations that lead to biochemically similar amino acids with homogeneous properties may not bear such strong selective advantages and could be regarded as almost neutral, while the mutations giving rise to extremely dissimilar amino acids may be deleterious.

We set up “Complex” models based on empirical observations to determine whether a mutation would be neutral, beneficial, or deleterious. There are a few nonsynonymous substitutions predicted to have been fixed by positive selection in response to evolutionary conflict with supporting functional evidence. Demogines et al. (2013) identified adaptively evolving sites in the transferrin receptor gene *TfR1* in wild rodents to include amino acids R, K, N, I, and T, which corresponds with pairwise Miyata scores ranging from 0.4 to 3.37. Likewise, Charron et al. (2008) proposed sites in the plant gene eIF4E to be in an arms race conflict with viral proteins, which includes those with amino acids L, P, and A that give pairwise Miyata scores ranging from 0.06 to 2.76. We found that the Miyata scores for all proposed positively selected residues in these studies ranged from 0.05 to 3.37, with the majority of scores falling between 1.5 and 2.5.

With the above proposition, we first parameterized the selection schemes in the complex Conflict model based on the empirical observations, and then adopted the same selection schemes in the complex Buffering model to represent one biologically plausible possibility for the *Wolbachia*-*bam* interaction. In the complex Conflict model, when *Wolbachia* is present, nonsynonymous mutations that give rise to biochemically similar amino acids (0 < MS(AA_cur_, AA_mut_) $\leq$ 1) are regarded as neutral with selection coefficients *s* = 0 (fig S1, top); mutations leading to mildly different amino acids (1 < MS(AA_cur_, AA_mut_) $\leq$ 3) are considered beneficial with selection coefficients *s* > 0 (fig S1, top); and mutations become deleterious with selection coefficients *s* < 0 when they generate extremely dissimilar amino acids (MS(AA_cur_, AA_mut_) > 3) (fig S1, top), as they are likely to disrupt the biological function of *bam*. These cutoffs are consistent with the range of Miyata scores found at sites that are proposed to be adaptively evolving in response to an evolutionary conflict. In the *Wolbachia*-absence phase in the complex Conflict model, to preserve the current amino acid sequences, we still assume that mutations leading to similar amino acid changes (0 < MS(AA_cur_, AA_mut_) $\leq$ 1) are considered neutral, but any mutation that causes a dissimilar amino acid change (MS(AA_cur_, AA_mut_) > 1) is deleterious with a selection coefficient *s* < 0 (fig S1, top).

When *Wolbachia* is present in the complex Buffering model, any mutation that gives rise to a mildly biochemically different amino acid from the ancestral state (0 < MS(AA_anc_, AA_mut_) $\leq$ 3) is regarded as neutral with selection coefficients *s* = 0 (fig. S1, bottom) due to the protection by *Wolbachia*. However, mutations are considered deleterious with selection coefficients *s* < 0 when they generate extremely dissimilar amino acids (MS(AA_anc_, AA_mut_) > 3) (fig. S1, bottom), since they are likely to disrupt the biological function of *bam*. When *Wolbachia* is lost from the population, only mutations that converge back towards the biochemical characteristics of the initial ancestral state relative to the current amino acid are favored (MS(AA_anc_, AA_mut_) – MS(AA_anc_, AA_mut_) < -1) with a selection coefficient *s* > 0, while the more divergent mutations (MS(AA_anc_, AA_mut_) – MS(AA_anc_, AA_cur_) > 1) are deleterious with a selection coefficient *s* < 0(fig. 2, bottom). Any mutation in between (-1 $\leq$ MS(AA_anc_, AA_mut_) – MS(AA_anc_, AA_cur_) $\leq$ 1) is considered neutral (*s* = 0) since it does not cause a radical functional change in the amino acid to increase or decrease the fitness of an individual.
