## Supplemental Figures for "A model of functionally buffered deleterious mutations can lead to signatures of positive selection"

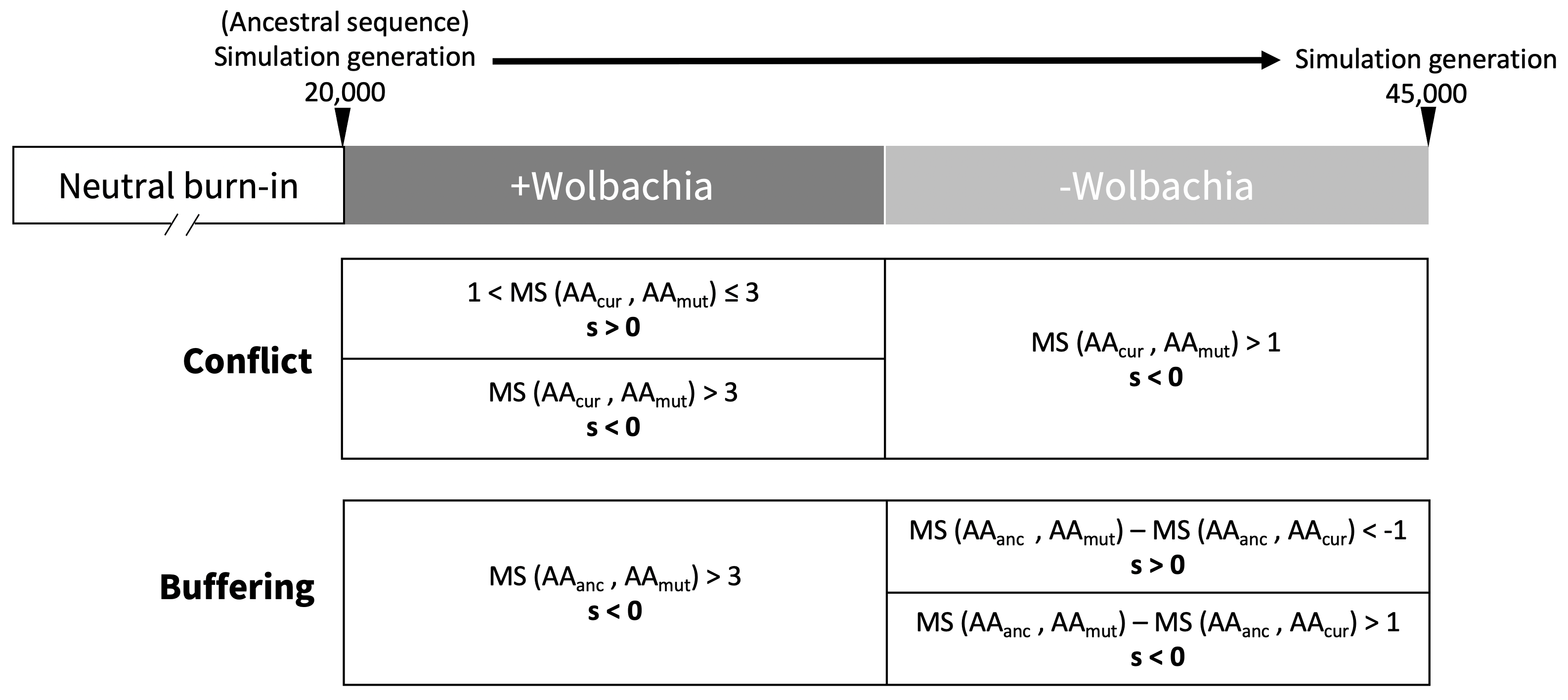

**Fig. S1. Simulation setup for Conflict and Buffering Complex models.** Selection on new nonsynonymous mutations (mutated amino acid, AA_mut_) is determined by their Miyata score (MS) to the appropriate reference amino acid (the current amino acid, AA_cur_ , or the ancestral amino acid, AA_anc_).

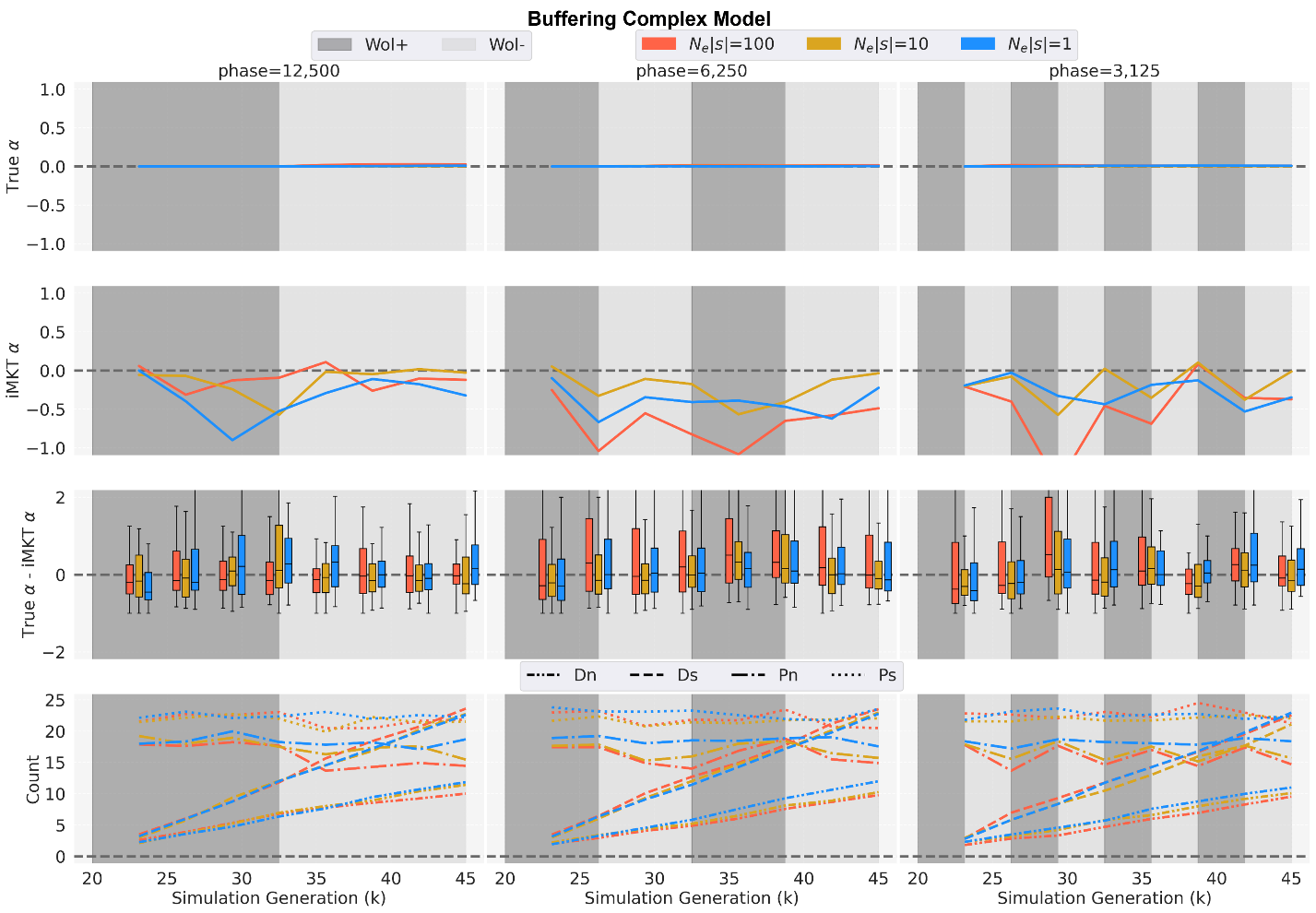

**Fig. S2. MK test results of simulations for Buffering Base and Complex models.** Each panel shows MK test analyses with different selection coefficients of *N_e_|s|*=100, *N_e_|s|*=10, and *N_e_|s|*=1 graphed across alternating phases (phase length=12,500, 6,250, and 3,125 simulation generations) of *Wolbachia* infection (Wol+, dark grey) and *Wolbachia* absence (Wol-, light grey) post-burn-in period. In each panel, row 1: the average true $\alpha$ in the simulations; row 2: the average estimated $\alpha$ (iMKT $\alpha$) in the simulations (FWW correction, SNPs frequency > 5% only); row 3: the distributions of differences between the true and estimated $\alpha$ every 3,125 simulation generations; row 4: The average of each MK test component (D_n_, D_s_, P_n_, P_s_).

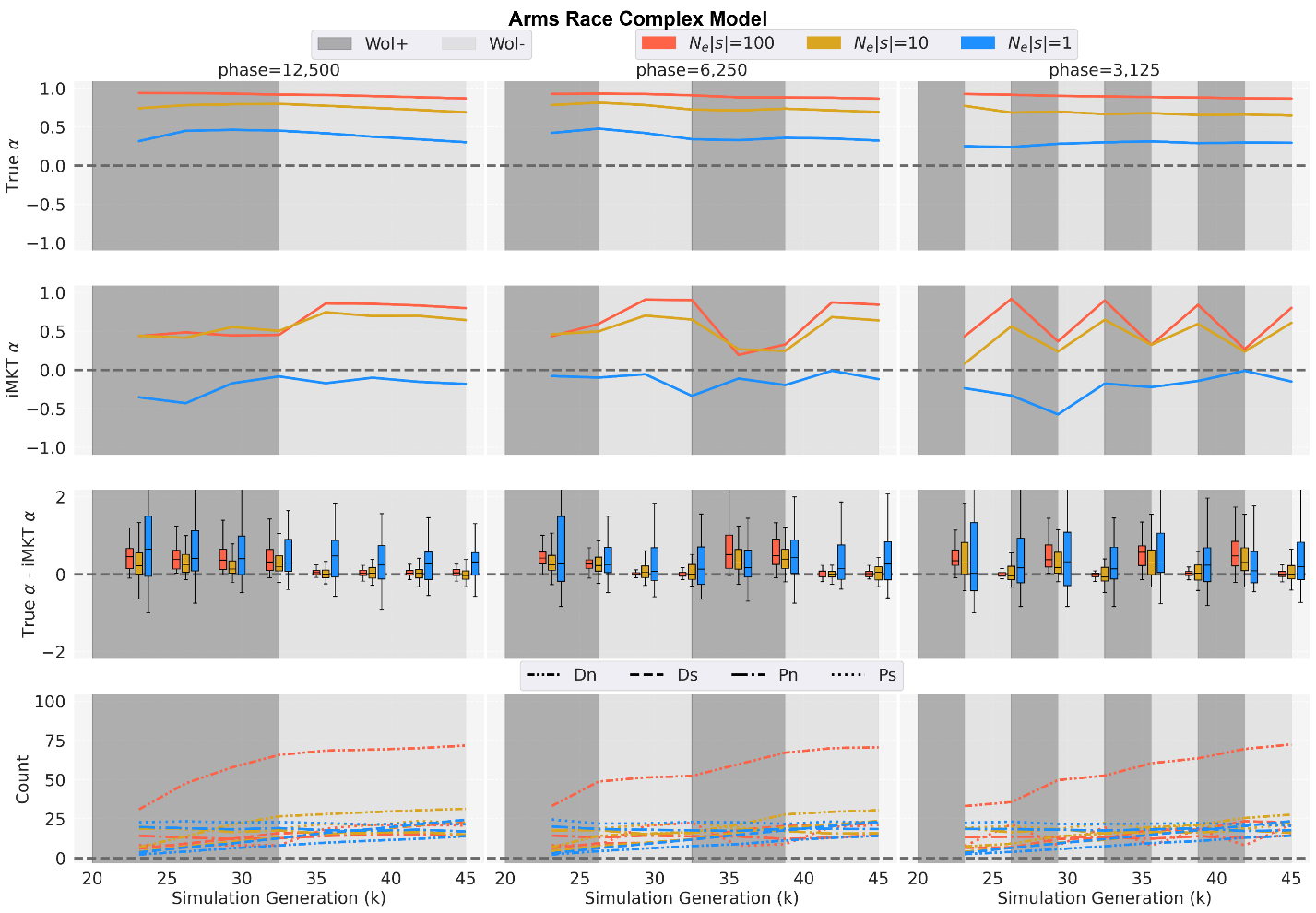

**Fig. S3.** **MK test results of simulations for Conflict Complex models:** Each panel shows MK test analyses with different selection coefficients of *N_e_|s|*=100, *N_e_|s|*=10, and *N_e_|s|*=1 graphed across alternating phases (phase length=12,500, 6,250, and 3,125 simulation generations) of *Wolbachia* infection (Wol+, dark grey) and *Wolbachia* absence (Wol-, light grey) post-burn-in period. In each panel, row 1: the average true $\alpha$ in the simulations; row 2: the average estimated $\alpha$ (iMKT $\alpha$) in the simulations (FWW correction, SNP frequency > 5% only); row 3: the distributions of differences between the true and estimated $\alpha$ every 3,125 simulation generations; row 4: The average of each MK test component (D_n_, D_s_, P_n_, P_s_).

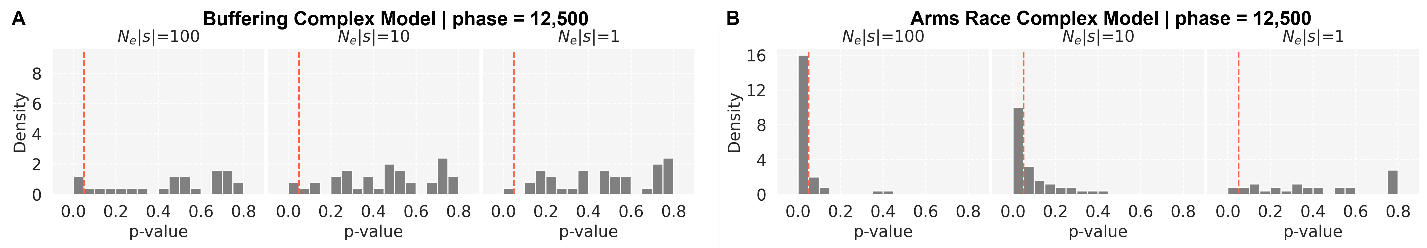
 **Fig. S4.** **Distributions of iMKT p-values.** iMKT p-values (FWW correction, SNPs frequency > 5% only) for simulation runs with *Wolbachia* phase=12,500 simulation generations and *N_e_|s|*=100, 10, 1 for the simulated models at 45,000 simulation generation. The vertical red line denotes p-value=0.05. Note that distributions are normalized to have an area of 1 under the histograms. (A) Buffering Complex model; (B) Conflict Complex model.

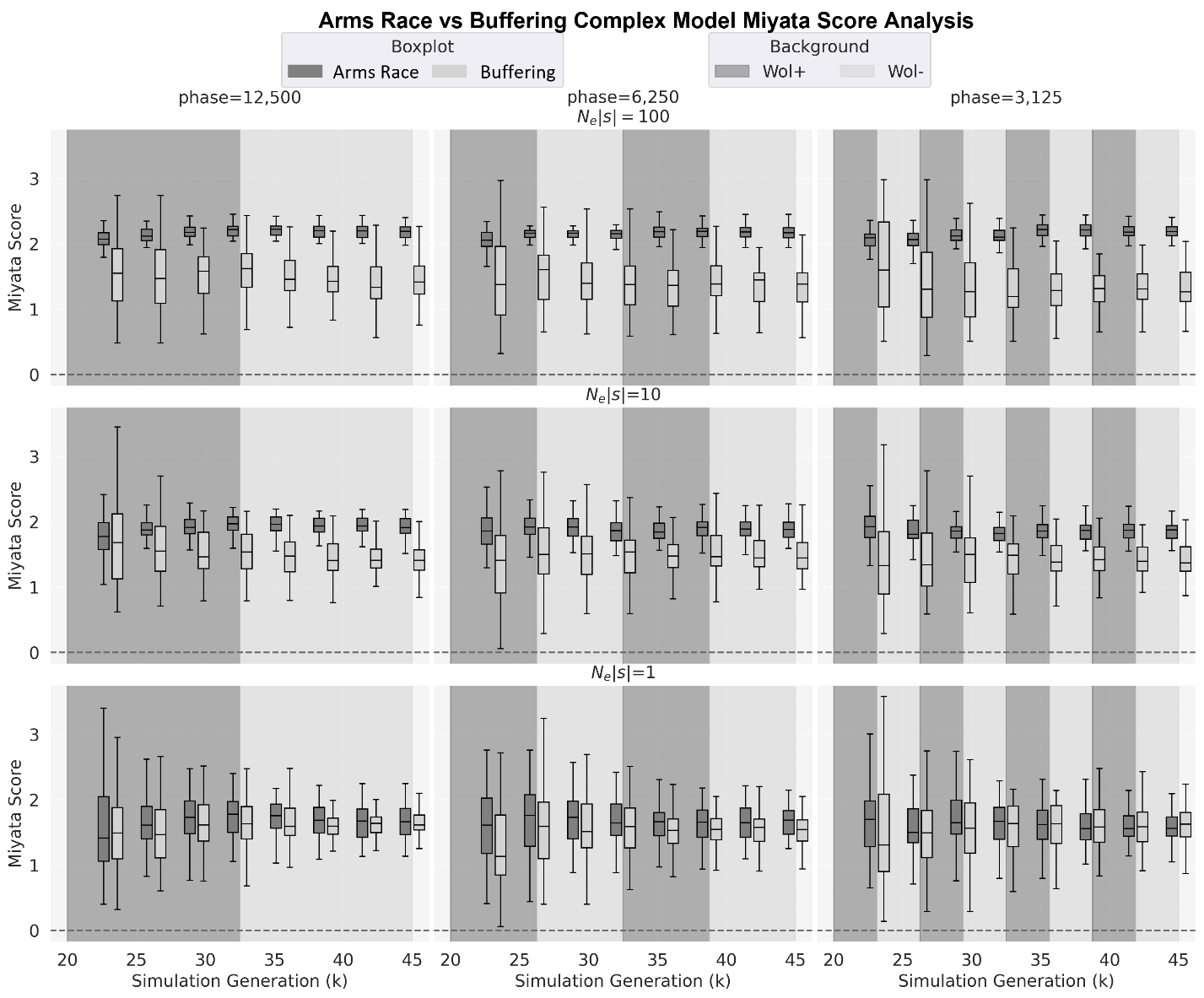

**Fig. S5. The distribution of Miyata scores per amino acid substitution for Complex models.** Miyata scores per amino acid substitution across multiple runs for substitutions between the consensus sequence at the end of a given simulation generation and the ancestral sequence. Data is shown for both the Conflict model (dark grey) and Buffering model (light grey) at every 3,125 simulated generations post burn-in for different phase lengths.

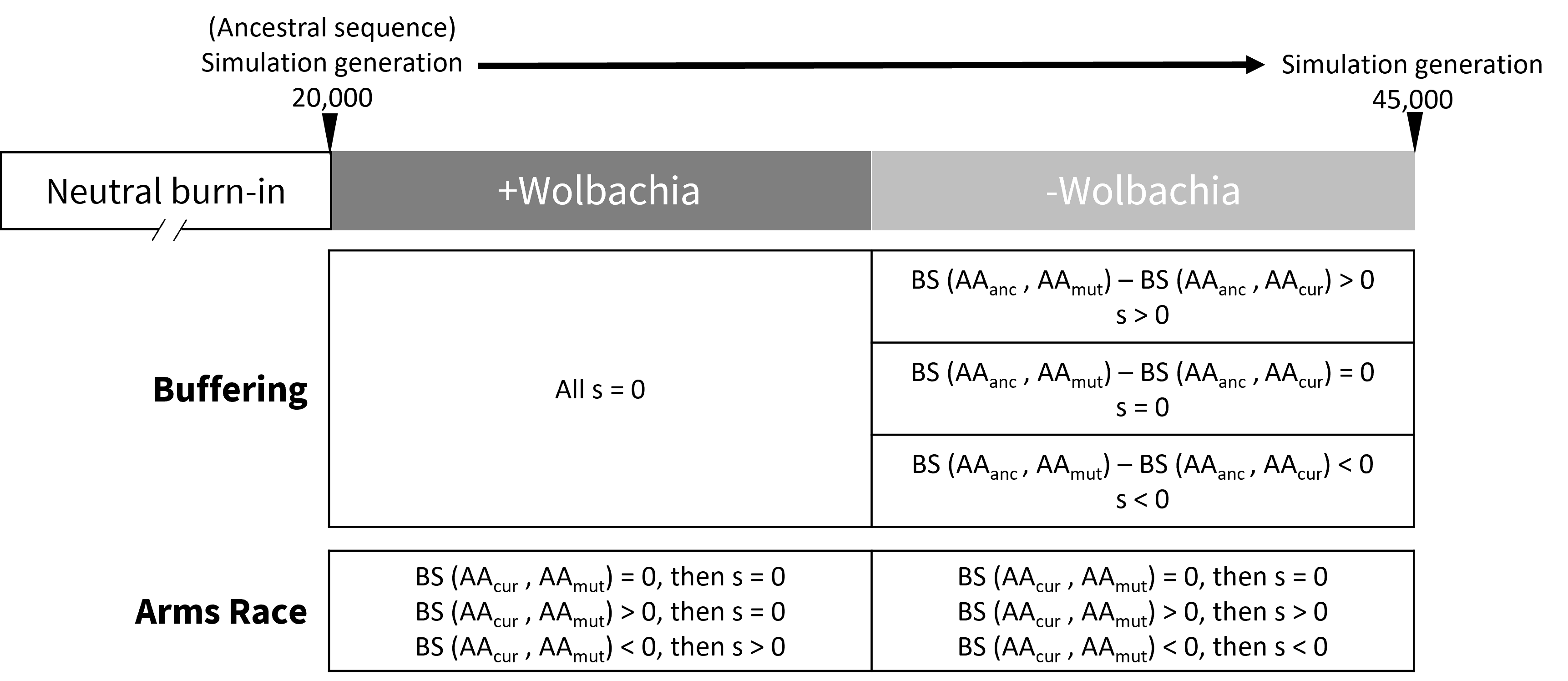
**Fig S6. Simulation setup for Arms Race and Buffering models using BLOSUM62 amino acid matrix for determining selection.** Selection on new nonsynonymous mutations (mutated amino acid, AA_mut_) is determined by their BLOSUM62 score (BS) to the appropriate reference amino acid (the current amino acid, AA_cur_ , or the ancestral amino acid, AA_anc_). BS = 0 represents an equal likelihood of finding one or the other amino acid at a site in the BLOSUM62 reference alignment; BS > 0 and BS < 0, respectively, represent a greater and lesser likelihood of finding the two amino acids aligned at the same site.

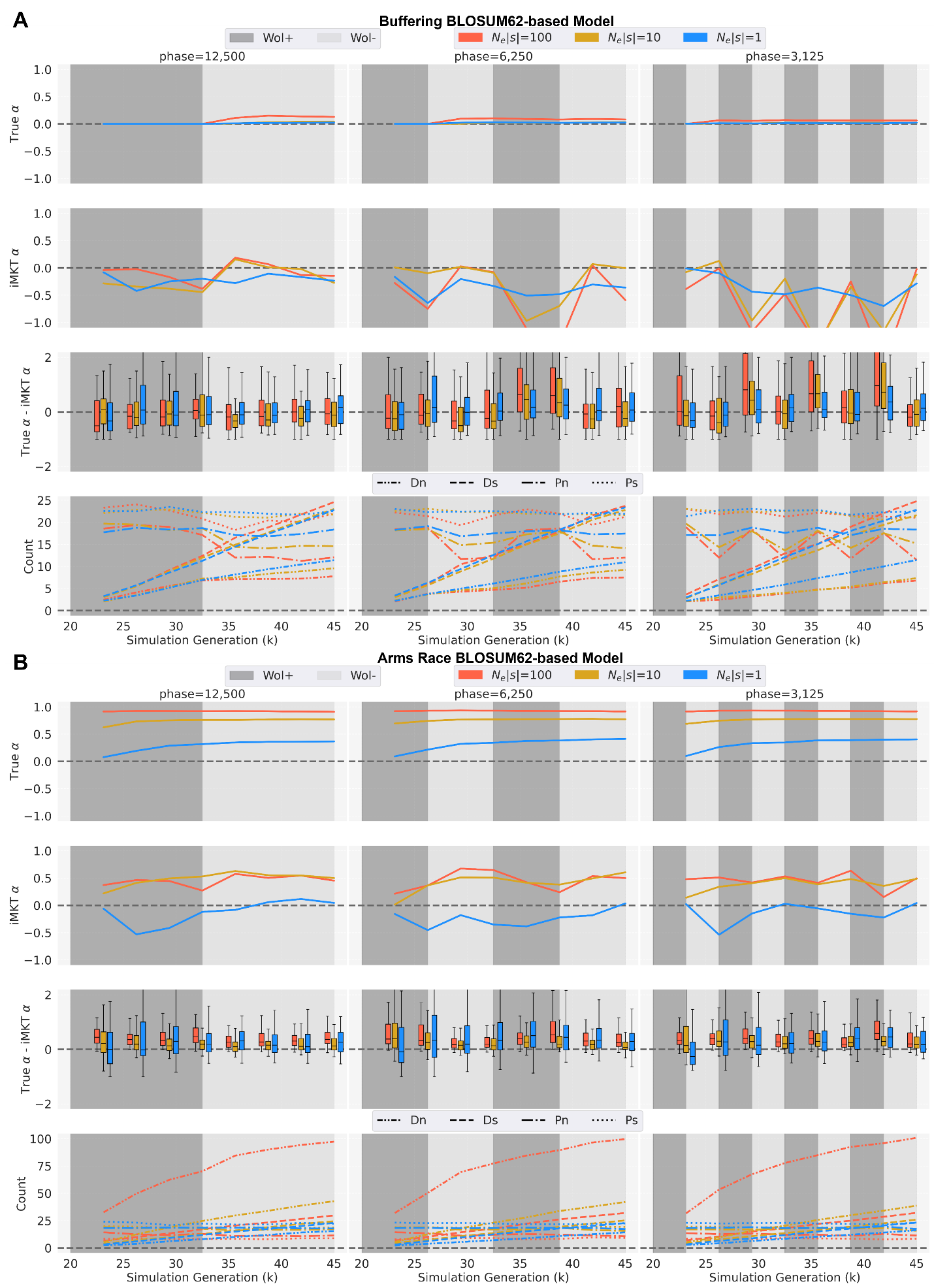

**Fig S7. MK test results from simulations using the BLOSUM62 amino acid matrix.** (A) Buffering model; (B) Arms Race model. Each panel shows MK test analyses with different selection coefficients of *N_e_|s|*=100, *N_e_|s|*=10, and *N_e_|s|*=1 graphed across alternating phases (phase length=12,500, 6,250, and 3,125 simulation generations) of *Wolbachia* infection (Wol+, dark grey) and *Wolbachia* absence (Wol-, light grey) post-burn-in period. In each panel, row 1: the average true $\alpha$ in the simulations; row 2: the average estimated $\alpha$ (iMKT $\alpha$) in the simulations (FWW correction, SNP frequency > 5% only); row 3: the distributions of differences between the true and estimated $\alpha$ every 3,125 simulation generations; row 4: The average of each MK test component (D_n_, D_s_, P_n_, P_s_).

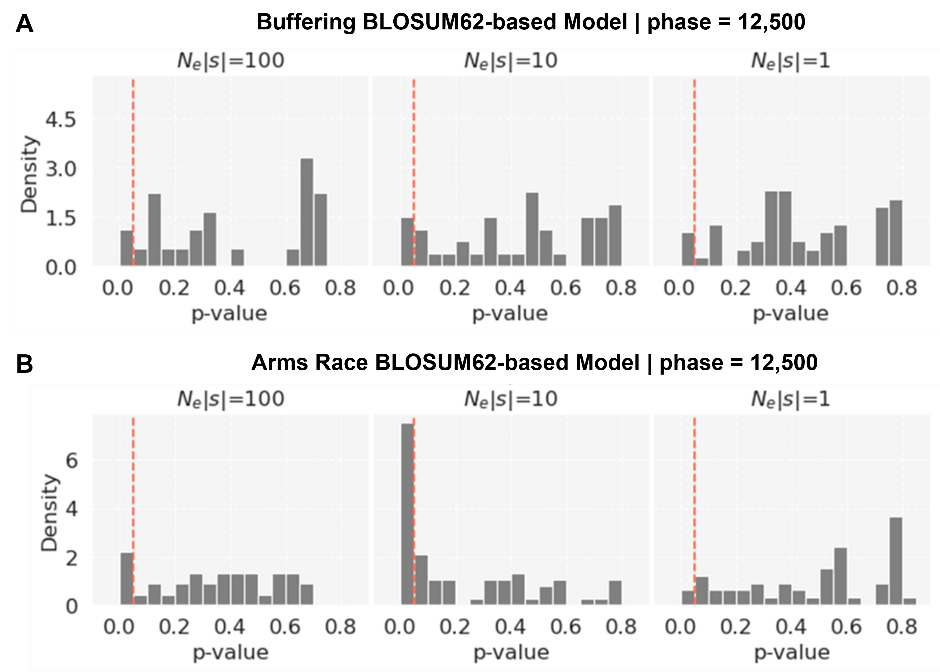

**Fig. S8. Distribution of MK test p-values from simulations using the BLOSUM62 amino acid matrix.** (A) Buffering Base model; (B) Arms Race Base model. MK test p-values (FWW correction, SNP frequency > 5% only) for simulation runs with *Wolbachia* phase=12,500 simulation generations and *N_e_|s|*=100, 10, and 1 for the simulated models at 45,000 simulation generations. The vertical red line denotes p-value=0.05. Distributions are normalized to have an area of 1 under the histograms.

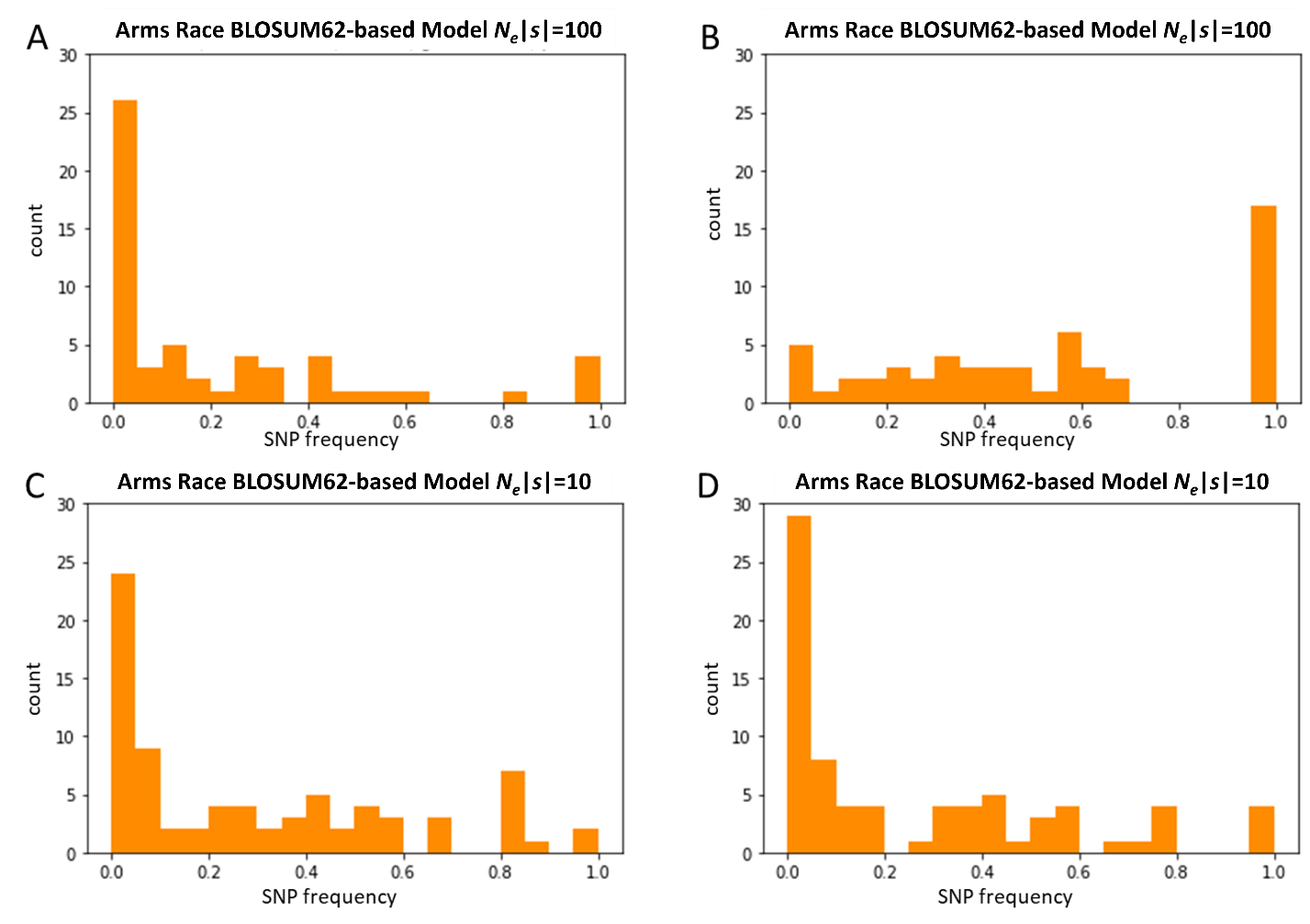
 **Fig. S9. Distribution of single nucleotide polymorphism (SNP) frequencies of simulations using the BLOSUM62 amino acid matrix.** Results are shown for *N_e_|s|*=100 and *N_e_|s|*=10, comparing (A and C) the distribution of all SNPs and (B and D) the distribution of only SNPs > 5% (FWW correction).

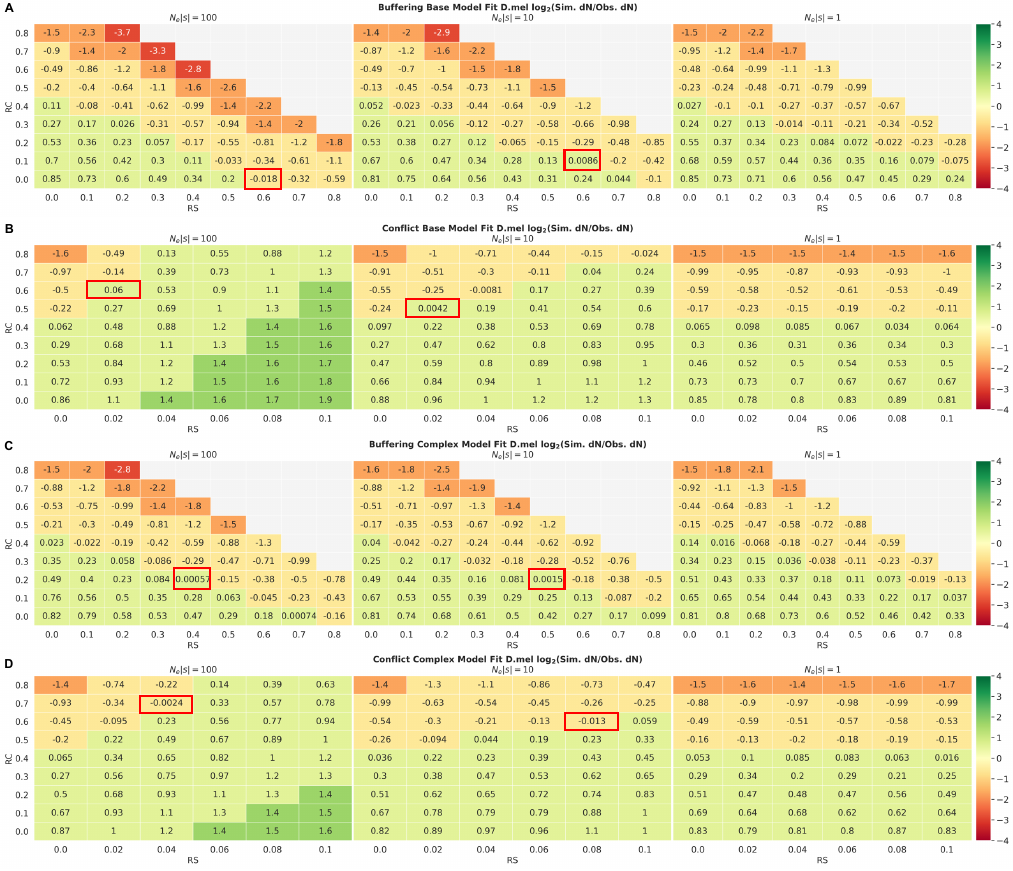

**Fig. S10. Heatmap of** $\boldsymbol{lo}\boldsymbol{g}_{\boldsymbol{2}}\frac{\boldsymbol{Simulation dN}}{\boldsymbol{Observed dN}}$ **for four models to fit the empirical data for Base and Complex models.** For each model, average dN was calculated across 50 simulation runs per each pair of conserved-site ratio and selection-site ratio to find the combination of parameters best fitting the empirical dN. Red boxes highlight the pairs of parameters used to investigate which model is preferred to recapitulate the observed data in MK test results and Miyata score analysis.

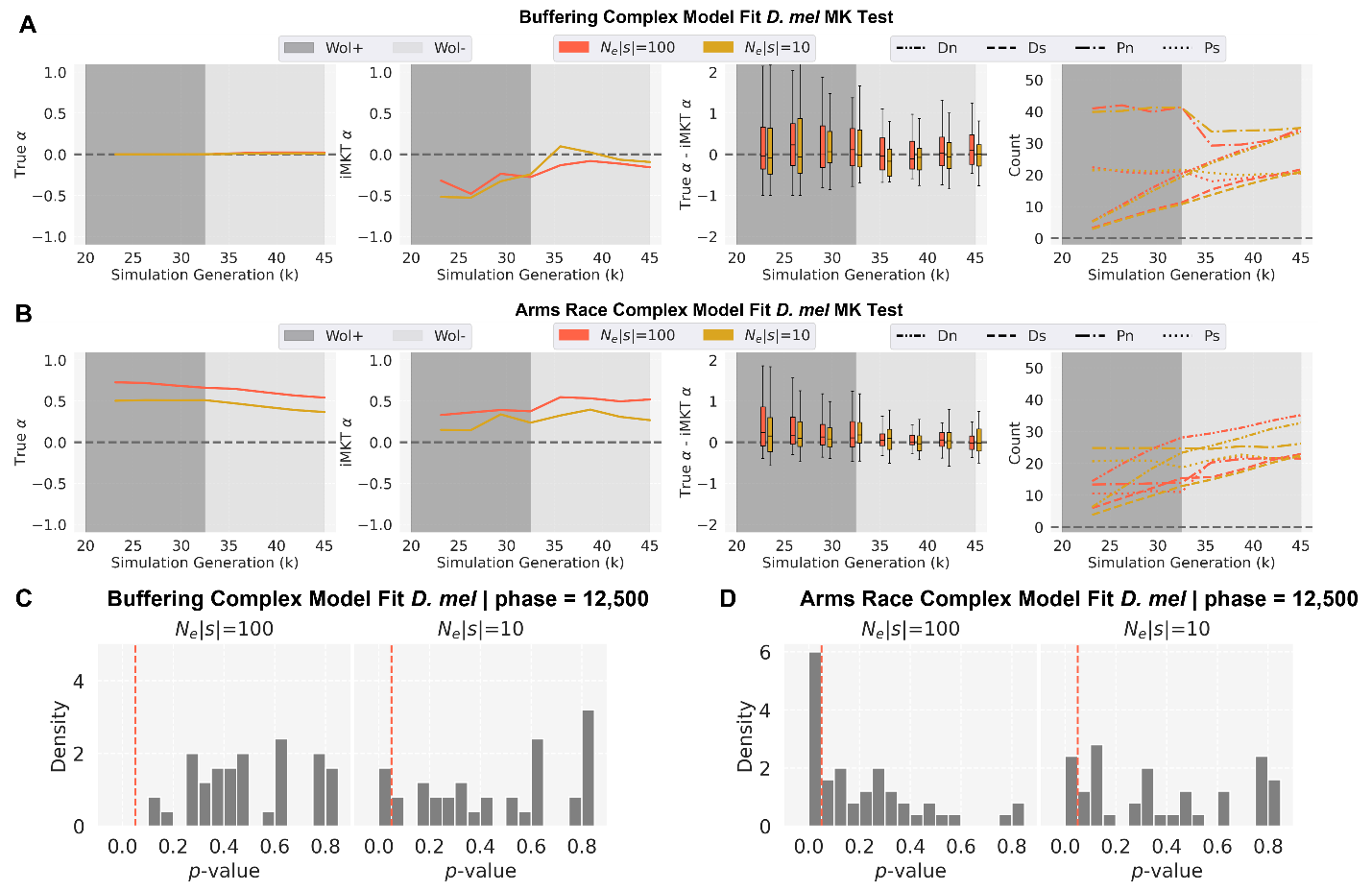

**Fig. S11. MK test results of simulations for best-fit Complex models for *D. melanogaster*.** (A-B) MK test $\alpha$ analysis of each model with different selection coefficients of *N_e_|s|*=100, *N_e_|s|*=10 graphed at phase length=12,500 simulation generations of *Wolbachia* infection (Wol+, dark grey) and *Wolbachia* absence (Wol-, light grey) post-burn-in period. In each panel, column 1: the average true $\alpha$ in the simulations; column 2: the average estimated $\alpha$ (iMKT $\alpha$)in the simulations (FWW correction, SNP frequency > 5% only); column 3: the distributions of differences between the true and estimated $\alpha$ every 3,125 simulation generations; column 4: The average of each MK test component (D_n_, D_s_, P_n_, P_s_). (C-D) Distributions of MK test p-values (FWW correction, SNPs frequency > 5% only) for simulation runs with *Wolbachia* phase=12,500 simulation generations and *N_e_|s|*=100 and 10 for the simulated models at 45,000 simulation generation. The vertical red line denotes p-value=0.05. Note that distributions are normalized to have an area of 1 under the histograms.

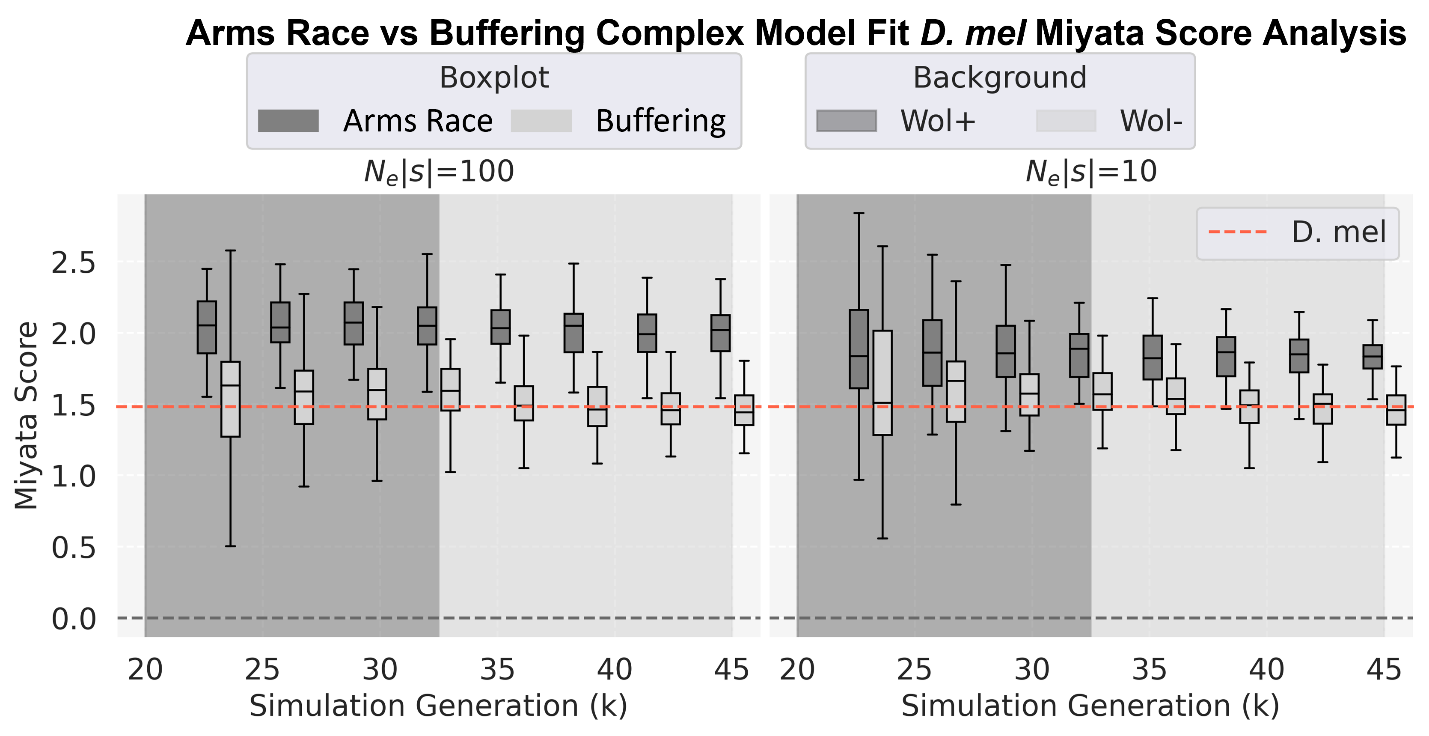
**Fig. S12. Distributions of Miyata score for the *D. melanogaster* samples from the simulations with best-fit parameters for the Complex models.** The boxplots are the distributions of Miyata scores for each model at every 3,125 generation. The distributions are compared with the observed summary statistics of *D. melanogaster* empirical data (red horizontal line).

| Type | 1 | 2 | 3 | 4 |
| --- | --- | --- | --- | --- |
| *D. melanogaster* | R | P | G | - |
| *D. simulans* | R | P (63) and S (5) | A | F |
| Predicted ancestral | R | P | A | F |
| Assigned status | Identical | Identical | Not identical | Not identical |

**Table S1.** Classification approach for assigning conserved and not conserved status to each position in the alignment of *D. melanogaster* and *D. simulans*. Conserved sites are assigned the selection coefficient *s* = -0.1 in the simulations. Type 1-4 show the four types of amino acid sites that were considered. Type 1 sites were classified as identical (possibly conserved) amino acids in both species. Type 2 sites were also classified as identical when polymorphic in one species with the major allele identical to that in the other. Type 3 has a substitution and Type 4 has an indel, with both being thus classified as not identical (not conserved) sites.
